# STAT3 protects HSCs from intrinsic interferon signaling and loss of long-term blood-forming activity

**DOI:** 10.1101/2023.02.10.528069

**Authors:** Bhakti Patel, Yifan Zhou, Rachel L. Babcock, Feiyang Ma, Malgorzata A. Zal, Dhiraj Kumar, Yusra B. Medik, Laura M. Kahn, Josué E. Pineda, Elizabeth M. Park, Ximing Tang, Maria Gabriela Raso, Tomasz Zal, Karen Clise-Dwyer, Filippo G. Giancotti, Simona Colla, Stephanie S. Watowich

## Abstract

STAT3 function in hematopoietic stem and progenitor cells (HSPCs) has been difficult to discern as *Stat3* deficiency in the hematopoietic system induces systemic inflammation, which can impact HSPC activity. To address this, we established mixed bone marrow (BM) chimeric mice with CreER-mediated *Stat3* deletion in 20% of the hematopoietic compartment. *Stat3*-deficient HSPCs had impaired hematopoietic activity and failed to undergo expansion in BM in contrast to *Stat3*-sufficient (CreER) controls. Single-cell RNA sequencing of Lin^−^ckit^+^Sca1^+^ BM cells revealed altered transcriptional responses in *Stat3*-deficient hematopoietic stem cells (HSCs) and multipotent progenitors, including intrinsic activation of cell cycle, stress response, and interferon signaling pathways. Consistent with their deregulation, *Stat3*-deficient Lin^−^ckit^+^Sca1^+^ cells accumulated γH2AX over time. Following secondary BM transplantation, *Stat3*-deficient HSPCs failed to reconstitute peripheral blood effectively, indicating a severe functional defect in the HSC compartment. Our results reveal essential roles for STAT3 in HSCs and suggest the potential for using targeted synthetic lethal approaches with STAT3 inhibition to remove defective or diseased HSPCs.

**Key Points:**

- STAT3 is critical for hematopoietic activity and hematopoietic stem cell maintenance in non-inflammatory conditions
- STAT3 has a cell-intrinsic role in the suppression of interferon signaling and myeloid-skewed transcription in hematopoietic stem cells

## Introduction

In homeostatic conditions, the replenishment of mature immune cells is sustained by relatively quiescent hematopoietic stem cells (HSCs), which undergo differentiation to generate mature blood lineages or self-renew to maintain an adequate stem cell pool (Laurenti and Gottgens, 2018; Orkin and Zon, 2008; Wilson et al., 2008). The hematopoietic system is highly dynamic and can adapt to physiological stress by increasing blood cell production. For instance, HSCs respond rapidly to molecular cues that indicate pathogen invasion or an increased demand for peripheral blood (PB) populations; such cues trigger enhanced HSC proliferation and differentiation, resulting in elevated numbers of mature hematopoietic cells in the periphery (Gomes et al., 2016; Orford and Scadden, 2008; Pinho and Frenette, 2019; Wei and Frenette, 2018; Zhao and Baltimore, 2015). Following the resolution of a transient demand, HSCs reestablish homeostasis and return to a largely dormant state (Pietras et al., 2011; Wilson et al., 2008). However, the dysregulation of these mechanisms can result in persistent HSC activation and dysfunction, subsequently increasing the propensity for bone marrow (BM) failure, preleukemic transformation, or inflammatory disorders (Yamashita et al., 2020; Zon, 2008). Identifying the intrinsic mechanisms that regulate HSCs is crucial to improving our understanding of hematopoiesis and devising therapeutic interventions for hematopoietic disorders (Chavakis et al., 2019; Pietras, 2017).

Ample evidence indicates that chronic inflammatory signals from cytokines, infectious microbes, growth factors, or Toll-like receptor (TLR) agonists can trigger the loss of HSC quiescence as well as HSC exhaustion, age-associated hematopoietic deregulation (inflammaging), and the impairment of long-term HSC activity (Bogeska et al., 2022; Chavakis et al., 2019; Esplin et al., 2011; Higa et al., 2021; Hormaechea-Agulla et al., 2020; King and Goodell, 2011; Kovtonyuk et al., 2022; Matatall et al., 2016; Muto et al., 2020; Pietras et al., 2016; Trowbridge and Starczynowski, 2021). Excessive exposure to these factors can deregulate HSCs directly or through changes in key components of the HSC niche (Chen et al., 2015; Essers et al., 2009; McCabe et al., 2015; Qin et al., 2017). For instance, chronic TLR signaling via NF-κB or sustained exposure to interleukin-1 degrades HSC function and leads to myeloid-skewed hematopoiesis (Boiko and Borghesi, 2012; Chavakis et al., 2019; Esplin et al., 2011; Liu et al., 2015; Pietras et al., 2016). Furthermore, the deregulation of intrinsic anti-inflammatory factors in HSCs, such as miR-146a, can permit the excessive activation of inflammatory signaling cascades, leading to HSC damage and BM failure (Starczynowski et al., 2010; 2011).

The signal transducer and activator of transcription 3 (STAT3) is a cytokine-responsive transcriptional regulator involved in numerous immune and hematopoietic functions. STAT3 has a significant anti-inflammatory role in mature myeloid cells, dendritic cells (DCs), and non-immune cell populations (Hillmer et al., 2016; Takeda et al., 1999; Williams et al., 2007). For instance, STAT3 inhibits NF-κB–mediated pro-inflammatory cytokine production in BM-derived macrophages and suppresses interferon (IFN) signaling in type I conventional DCs (Chrisikos et al., 2022; Zhang et al., 2014). However, STAT3’s anti-inflammatory function has prevented us from obtaining an understanding of its intrinsic role in HSCs, as *Stat3* deletion in the full hematopoietic compartment or in myeloid cells or DCs specifically leads to systemic inflammation (Mellilo et al., 2010; Panopoulos et al., 2006; Takeda et al., 1999; Welte et al., 2003; Zhang et al., 2018). Previous studies implicated STAT3 in the regulation of HSCs yet these were confounded by the systemic inflammation that accompanies *Stat3* deletion, or by the transplantation of adult BM cells with constitutively activated STAT3 (Chung et al., 2006; Hong et al., 2014; Mantel et al., 2012; Oh and Eaves, 2002; Zhang et al., 2018). Thus, novel approaches are needed to delineate the physiological role of STAT3 in HSCs in the absence of systemic inflammation.

Here, we used competitive mixed BM chimeric mice with conditional *Stat3* deletion in approximately 20% of the hematopoietic compartment to establish a model of *Stat3*-deficiency in the hematopoietic stem and progenitor (HSPC) compartment on a non-inflammatory background. We found that STAT3 is required for effective hematopoiesis and long-term HSC function in the absence of peripheral inflammation. This role is mediated by the STAT3-dependent control of HSC cell cycle regulation, prevention of intrinsic IFN signaling, and restraint of myeloid-skewed transcription.

## Results

### STAT3 is required for hematopoietic activity in the absence of peripheral inflammation

To investigate the roles of STAT3 in HSPCs in non-inflammatory conditions, we established mixed BM chimeric mice with conditional *Stat3* deletion in approximately 20% of the hematopoietic compartment (Fig. 1 A). Our approach enabled us to track CD45.2^+^ CreER *Stat3*^f/f^ (*Stat3*-deficient) or CreER (*Stat3*-sufficient control) cells and CD45.1^+^CD45.2^+^ wild-type (WT) competitors in the same background. After confirming engraftment, we treated mice with tamoxifen to induce *Stat3* deletion (Fig. 1 A). An analysis of circulating cytokines and chemokines and colon histopathology revealed negligible differences between mice with CreER *Stat3*^f/f^ or CreER BM 8 weeks after tamoxifen treatment, and effective *Stat3* depletion was confirmed in CD45.2^+^ CreER *Stat3*^f/f^ BM (Fig. 1 B-D, Table S1, Table S2, and Table S3). These data indicate that our mixed BM chimeric mice enable the evaluation of STAT3’s function in non-inflammatory conditions.

**Figure 1.**
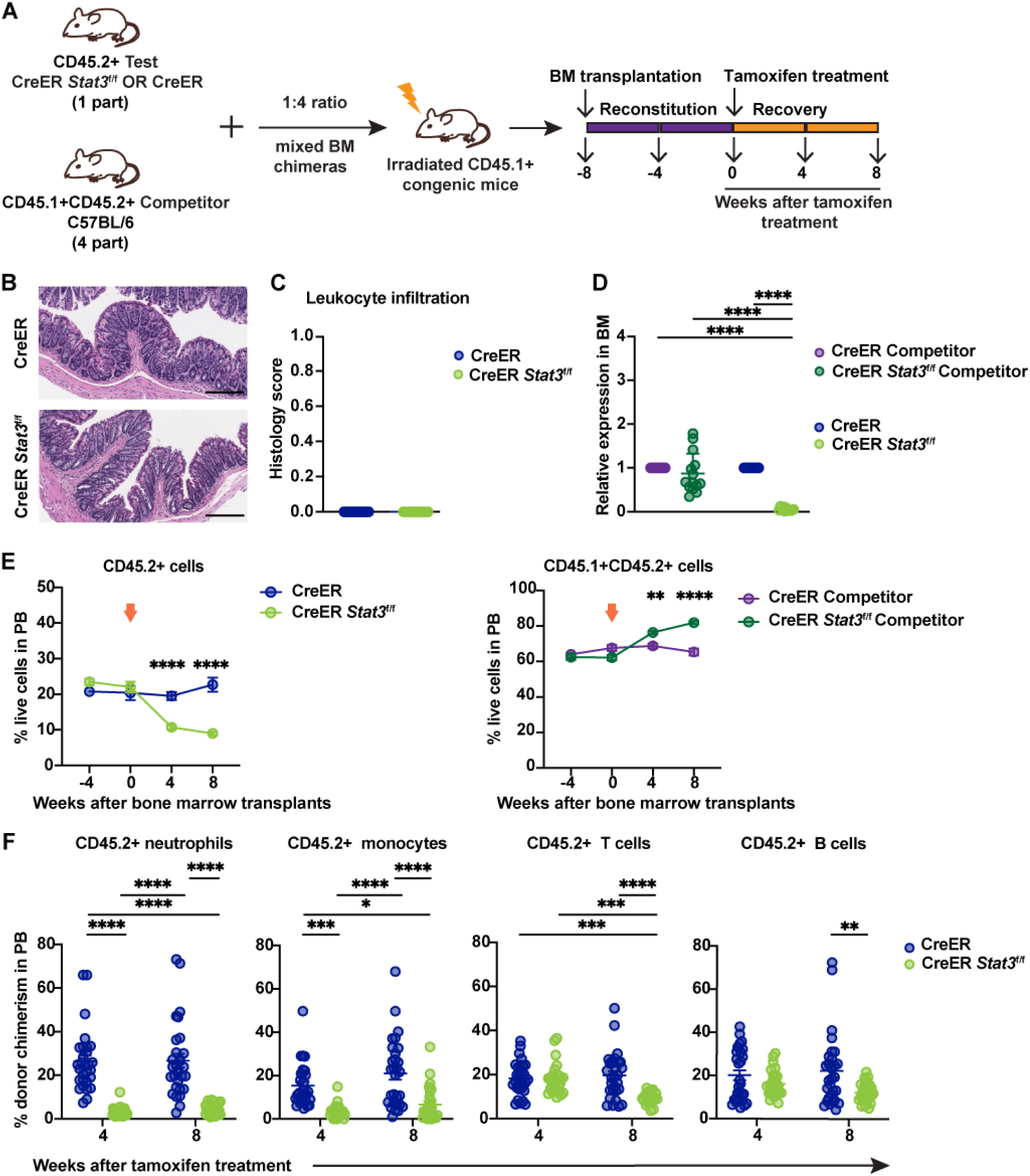
*Stat3*-deficient HSPCs show ineffective hematopoietic activity in non-inflammatory conditions. (A) A schematic diagram illustrating the mixed BM chimeric model system. (B and C) A trained veterinary pathologist performed histological assessments of colon tissues from BM chimeric mice with CreER (*Stat3*-sufficient) or CreER *Stat3*^f/f^ (*Stat3*-deficient) cells 8 weeks after tamoxifen treatment (n = 11-14 mice per group). Representative photomicrographs of H&E-stained colon tissue sections. Scale bars represent 200 μm (B). Quantification of indicated histology feature of the H&E-stained colon tissue sections (C). (D) qRT-PCR analysis of *Stat3* expression in CD45.2^+^ test and CD45.1^+^CD45.2^+^ WT competitor cells (n = 11-14 mice per group). (E) Frequencies of CD45.2^+^ test and CD45.1^+^CD45.2^+^ WT competitor cells in the PB of chimeric mice at the indicated times before and after (arrows) tamoxifen treatment. (F) Donor chimerism of CD45.2^+^ neutrophils (CD11b^+^Ly6G^+^Ly6C^low^), monocytes (CD11b^+^Ly6G^−^Ly6C^high^), T cells (CD3^+^), and B cells (B220^+^) in the PB of chimeric mice at the indicated times after tamoxifen treatment. Donor chimerism was determined as the percentage of the indicated population within the combined CD45.1^+^, CD45.2^+^, and CD45.1^+^CD45.2^+^ cells of the same population. Data in E and F are representative of 2 independent experiments (n = 26-29 mice per group). Error bars indicate means ± SEMs. Statistical analyses were performed using 2-tailed unpaired Student *t*-test (C), 2-way ANOVA with the Sidak (E) or Tukey (D, F) multiple comparison test. \**P* < 0.05; \*\**P* < 0.01; \*\*\**P* < 0.001; \*\*\*\**P* < 0.0001.

The ability of CreER *Stat3*^f/f^ HSPCs to generate PB was significantly impaired 4 weeks (*P* < 0.001) and 8 weeks (*P* < 0.0001) after *Stat3* deletion, as indicated by the loss of circulating CD45.2^+^ cells (Fig. 1 E, left panel). This loss was accompanied by an increase in WT competitor cells (Fig. 1 E, right panel), suggesting a competitive disadvantage for *Stat3*-deficient HSPCs. Furthermore, CreER *Stat3*^f/f^ myeloid cells, neutrophils, monocytes, and T and B cells in PB were significantly depleted 8 weeks after *Stat3* deletion (Fig. 1 F and Fig. S1 A), whereas circulating WT competitor populations in mice with *Stat3* deletion showed increased chimerism (Fig. S1, B and C). We also observed a significant loss of CreER *Stat3*^f/f^ immune subsets in spleen and colonic lamina propria, while WT competitor populations were increased in spleens of mice with *Stat3* deletion (Fig. 2 and Fig. S1, D-H). These data indicate that STAT3 is crucial for sustaining the effective production of mature hematopoietic cells in the absence of peripheral inflammation.

**Figure 2.**
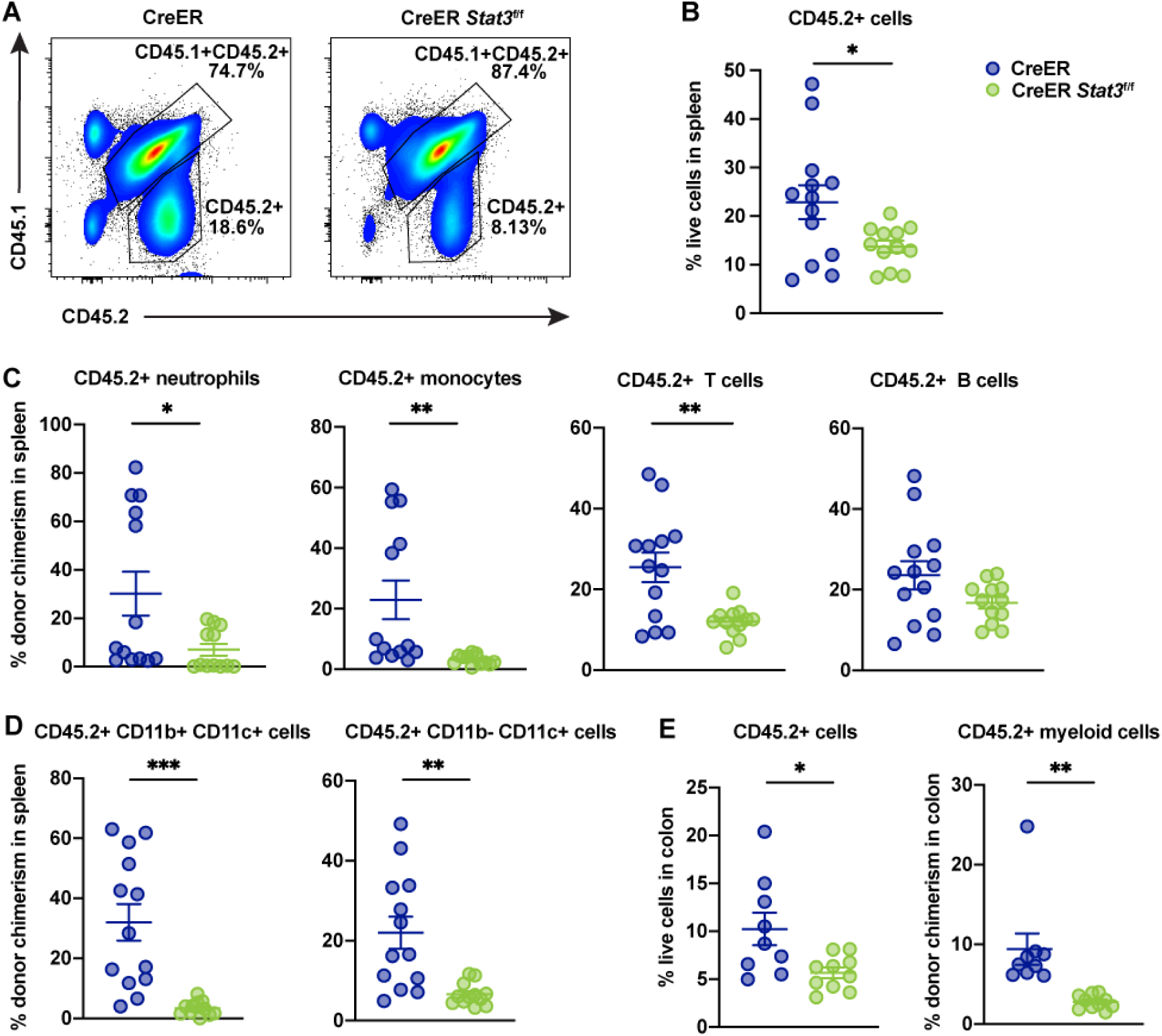
*Stat3* deletion in HSPCs results in the loss of peripheral immune cell lineages. Major immune cell lineages in the spleens and colons of BM chimeric mice with CreER (*Stat3*-sufficient) or CreER *Stat3*^f/f^ (*Stat3*-deficient) cells were evaluated 8 weeks after tamoxifen treatment. (A) Representative flow cytometric plots showing proportions of CD45.2^+^ test and CD45.1^+^CD45.2^+^ WT competitor cells in spleen. (B) Frequencies of CD45.2^+^ test cells in spleen (n = 12 or 13 mice per group). (C) Donor chimerism of CD45.2^+^ neutrophils (CD11b^+^Ly6G^+^Ly6C^low^), monocytes (CD11b^+^Ly6G^−^Ly6C^high^), T cells (CD3^+^), and B cells (B220^+^) in spleen. (D) Donor chimerism of CD45.2^+^ DC subsets (CD11b^+^CD11c^+^ and CD11b^−^ CD11c^+^) in spleen. (E) Frequencies of CD45.2^+^ cells (left) and donor chimerism of CD45.2^+^CD11b^+^ myeloid cells (right) in colon (n = 9 or 10 mice per group). Data in B-E are representative of 2 independent experiments. Error bars indicate means ± SEMs. Statistical analyses were performed using a 2-tailed unpaired Student *t*-test. \**P* < 0.05; \*\**P* < 0.01; \*\*\**P* < 0.001.

### STAT3 mediates the expansion of BM HSPCs

To further understand STAT3 function, we evaluated BM populations in chimeric mice 8 weeks after *Stat3* deletion. CD45.2^+^ cells, as well as CD45.2^+^ Lin^−^ cells, myeloid (CD11b^+^) cells, and Gr1^+^ neutrophils, were significantly reduced in CreER *Stat3*^f/f^ BM chimeras, whereas WT competitor populations were increased (Fig. 3, A-C). In CreER BM chimeras, CD45.2^+^ LSKs, long-term HSCs (LT-HSCs), myeloid-biased multipotent progenitors (MPP-Mye), lymphoid-biased MPPs (MPP-Ly), and committed progenitor subsets were increased, whereas CreER *Stat3*^f/f^ populations were not (Fig. 3 D and Fig S2, A-C). In addition, CreER BM chimeras showed reduction of WT competitor HSPCs after tamoxifen treatment (Fig. S2 D), while this was lessened in CreER *Stat3*^f/f^ chimeric mice (Fig. S2 D), suggesting distinct competitive effects in the presence of CreER or CreER *Stat3*^f/f^ HSPCs. Collectively, these data suggest that tamoxifen or CreER activation stimulates HSPC production, which is mediated by STAT3.

**Figure 3.**
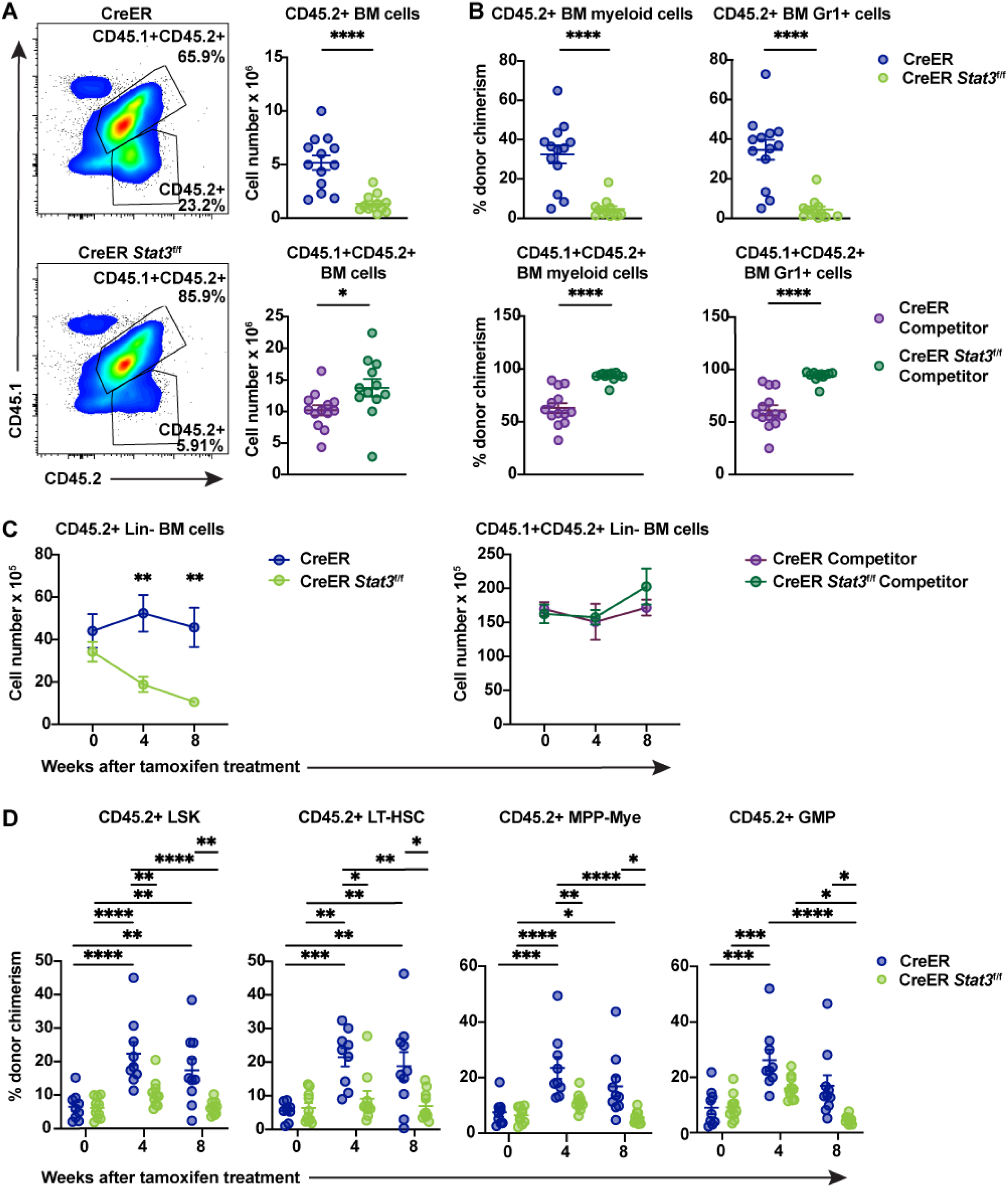
STAT3 is required to maintain BM HSPCs. Major immune cell lineages in total BM from chimeric mice were evaluated 8 weeks after tamoxifen treatment. (A) Representative flow cytometric plots of CD45.2^+^ test and CD45.1^+^CD45.2^+^ WT competitor cells (left) and absolute counts of these cells (right) in the BM of chimeric mice with CreER (*Stat3*-sufficient) or CreER *Stat3*^f/f^ (*Stat3*-deficient) BM. (B) Donor chimerism of CD45.2^+^ test (upper panels) and CD45.1^+^CD45.2^+^ WT competitor (lower panels) myeloid (CD11b^+^) and Gr1^+^ cells (n = 12 or 13 mice per group). (C and D) Lin^−^ cells from BM chimeric mice were evaluated at the indicated times after *Stat3* deletion (n = 9 or 10 mice per group). Absolute numbers of Lin^−^CD45.2^+^ test and CD45.1^+^CD45.2^+^ WT competitor cells in BM (C). Donor chimerism of CD45.2^+^ LSK (Lin^−^ ckit^+^Sca1^+^), LT-HSC (Lin^−^ckit^+^Sca1^+^CD135^−^CD34^−^CD48^−^CD150^+^), MPP-My (Lin^−^ ckit^+^Sca1^+^CD135^−^CD48^+^CD150^−^), and GMP (Lin^−^CD127^−^ckit^+^Sca1^−^CD16/32^+^CD34^+^) populations (D). Data are representative of 2 independent experiments. Error bars indicate means ± SEMs. Statistical analyses were performed using a 2-tailed unpaired Student *t*-test (A and B) or with 2-way ANOVA with the Sidak (C) or Tukey (D) multiple comparison test. \**P* < 0.05; \*\**P* < 0.01; \*\*\**P* < 0.001; \*\*\*\**P* < 0.0001.

To better understand the effects of tamoxifen and CreER activation on BM HSPCs, we treated WT C57Bl/6J and CreER mice with tamoxifen and assessed PB subsets and BM progenitors 4 weeks later (Fig. 4 A). WT mice treated with tamoxifen for <24 hours (i.e., at 0 weeks) served as the control group. With the exception of a modest reduction in neutrophil frequencies in WT mice, tamoxifen treatment and CreER activation did not affect the proportion of major PB lineages (Fig. 4, B and C). By contrast, the frequencies and absolute numbers of BM LT-HSCs were increased in WT mice 4 weeks after tamoxifen treatment (Fig. 4, D and E). Moreover, the percentage of BM LSKs and MPP-Mye were increased in CreER mice 4 weeks after tamoxifen treatment, compared with WT and CreER controls at 0 weeks or WT mice at 4 weeks. Common myeloid progenitor (CMP) frequencies were also increased by tamoxifen treatment in WT and CreER mice, whereas granulocyte-monocyte progenitors (GMPs), common lymphoid progenitors (CLPs), and megakaryocytic-erythroid progenitors (MEPs) remained unchanged (Fig. 4, D and E). These data suggest that tamoxifen stimulates the proliferation of LT-HSCs while the tamoxifen-mediated activation of CreER further affects LSK, MPP-Mye, and CMP populations. Together, our results suggest that STAT3 mediates the transient proliferation of LT-HSCs and the expansion of the HSPC compartment in response to tamoxifen and CreER activation.

**Figure 4.**
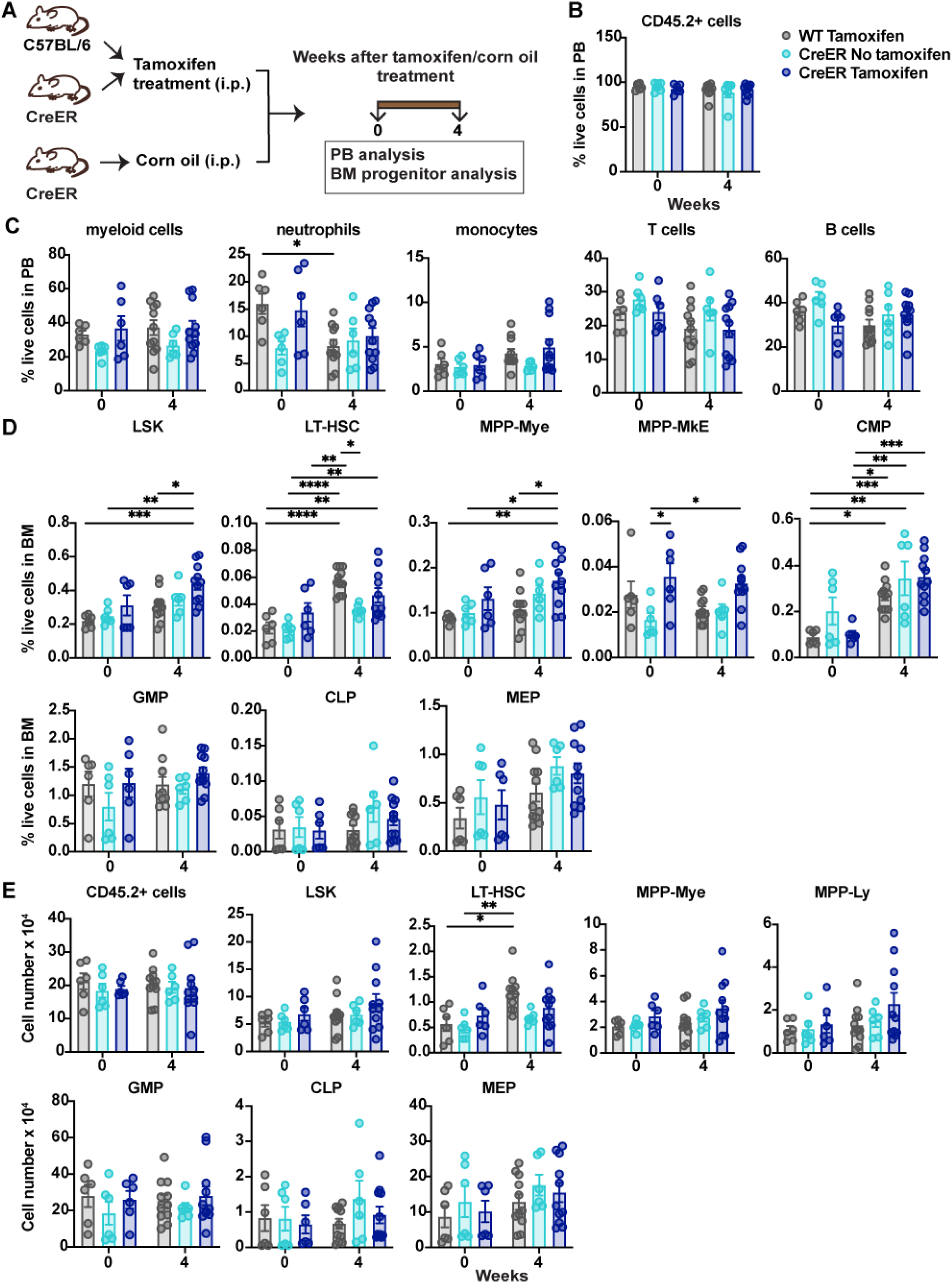
Evaluation of tamoxifen treatment in WT and CreER mice. (A) A schematic diagram showing the experimental design for analysis of WT or CreER mice treated with tamoxifen or corn oil (vehicle). (B-E) Major immune cell lineages in the PB and HSPC BM populations were evaluated 0 and 4 weeks after tamoxifen treatment. (B) Frequencies of CD45.2^+^ cells in PB. (C) Frequencies of mature immune cell lineages in PB. (D) Frequencies of the indicated HSPCs in the Lin^−^ BM population. (E) Absolute numbers of CD45.2^+^ cells in BM and absolute numbers of the indicated HSPCs in the Lin^−^ BM population. Data in B-E are representative of 2 independent experiments (n = 6-11 mice per group). Error bars indicate means ± SEMs. Statistical analyses were performed using 2-way ANOVA with the Tukey multiple comparison test. \**P* < 0.05; \*\**P* < 0.01; \*\*\**P* < 0.001; \*\*\*\**P* < 0.0001.

### STAT3 regulates HSPC transcriptional pathways

To elucidate the transcriptional responses mediated by STAT3 in HSPCs, we subjected LSKs purified from BM chimeric mice to single-cell RNA sequencing (scRNA-seq). For these assays, we used mice injected with a 1:1 ratio of CD45.2^+^ test (CreER or CreER *Stat3*^f/f^) and CD45.1^+^CD45.2^+^ WT competitor cells to ensure sufficient LSKs for sequencing. We confirmed a non-inflammatory background 8 weeks after *Stat3* deletion (Fig. 5 A and Fig. S3, A and B). We next purified LSKs by FACS and combined cells from individual mice for scRNA-seq: 12174 CD45.2^+^ CreER (*Stat3*-sufficient) LSKs and 9822 WT CD45.1^+^CD45.2^+^ LSK competitors from 15 CreER BM chimeric mice, and 8864 CD45.2^+^ CreER *Stat3*^f/f^ (*Stat3*-deficient) LSKs and 11322 WT CD45.1^+^CD45.2^+^ LSK competitors from 15 CreER *Stat3*^f/f^ BM chimeric mice.

**Figure 5.**
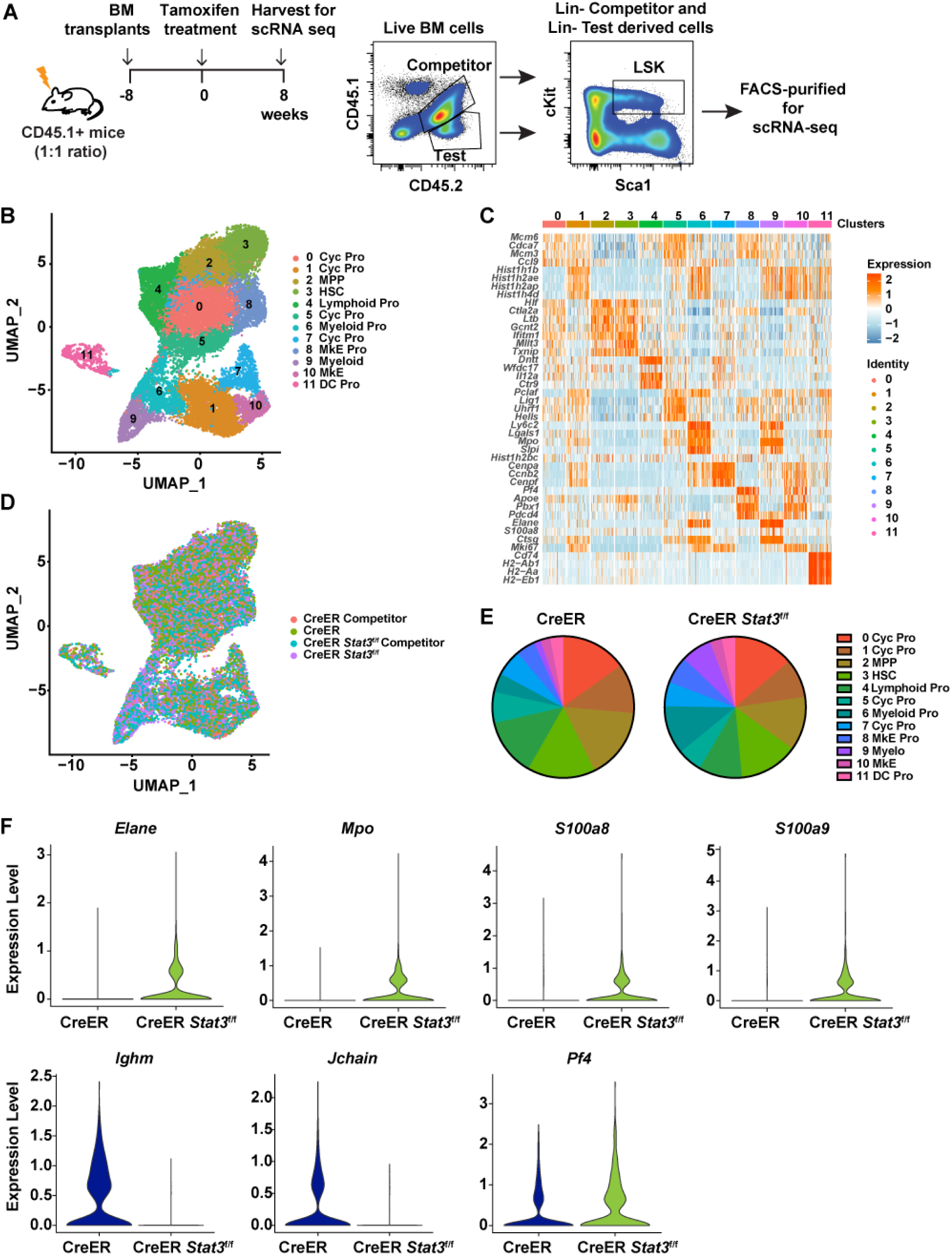
*Stat3-*deficient LSKs have a unique transcriptional profile. (A) A schematic diagram showing the experimental design for using 1:1 ratio of CD45.2^+^ test and CD45.1^+^CD45.2^+^ WT competitor cells to generate BM chimeric mice with sufficient LSKs for scRNA-seq analysis. (B) UMAP plot showing 12 clusters in the integrated LSK compartment, including 12174 CD45.2^+^ CreER LSKs and 9822 CD45.1^+^CD45.2^+^ WT competitor LSKs from CreER (*Stat3*-sufficient) BM chimeric mice and 8864 CD45.2^+^ CreER *Stat3*^f/f^ LSKs and 11322 CD45.1^+^CD45.2^+^ WT competitor LSKs from CreER *Stat3*^f/f^ (*Stat3*-deficient) BM chimeric mice. Results were pooled to generate a merged dataset (n = 15 mice per group). Different colors represent distinct clusters (or transcriptional states) based on differential gene expression signatures. Each dot represents a single cell. (C) Heatmap displaying cluster-defining stem cell– and lineage-specific gene expression. (D) UMAP plot showing the distribution of the 4 experimental groups in the integrated LSK compartment. Different colors represent different experimental groups. (E) Proportions of individual clusters represented as percentages of the total cells in each experimental group. (F) Violin plots showing the expression levels of myeloid lineage–specific genes (*Mpo, Elane, S100a8,* and *S100a9*), lymphoid-specific genes (*Ighm* and *Jchain*) and megakaryocyte-specific gene (*Pf4*) in cluster 3 (HSCs) from CreER LSKs and CreER *Stat3*^f/f^ LSKs.

Dimensionality reduction analysis of the integrated LSK compartment from all experimental groups revealed 12 distinct transcriptional states (Fig. 5 B). Individual clusters were defined based on the enriched expression of stem cell– and lineage-specific genes, revealing HSC- and MPP-like subpopulations as well as subpopulations defined by the expression of myeloid-, megakaryocyte/erythroid-, or lymphoid-specific genes (Fig. 5 C and Fig. S3 C). In all experimental groups, LSKs contributed to each transcriptionally-defined cluster; however, CreER *Stat3*^f/f^ LSKs showed enhanced accumulation in myeloid-biased (clusters 6 and 9) and megakaryocyte/erythroid (clusters 8 and 10) clusters (Fig. 5, D and E). Compared with CreER LSKs, CreER *Stat3*^f/f^ LSKs were less abundant in cycling progenitor clusters (clusters 1 and 5) and the lymphoid cluster (cluster 4) (Fig. 5, D and E). CreER *Stat3*^f/f^ HSCs and MPPs (clusters 3 and 2, respectively) had abundances similar to those of the CreER controls, yet these populations had increased expression of genes associated with myeloid (*Mpo*, *Elane*, *S100a8*, and *S100a9*) and megakaryocyte/erythroid (*Pf4*) lineages, and decreased expression of genes associated with lymphoid lineages (*Ighm* and *Jchain*) (Fig. 5 F and Fig. S3 D). These results suggest that STAT3 suppresses the overproduction of myeloid and megakaryocyte progenitors by restraining myeloid- and megakaryocyte-biased gene expression in developmentally early multipotent BM progenitors. These data also suggest that STAT3 controls the proliferative responses of HSPCs.

Pathway enrichment analysis revealed that, compared with the CreER control, the CreER *Stat3*^f/f^ HSC cluster had augmentation of multiple transcriptional pathways, including those involving Myc targets, IFN-α, oxidative phosphorylation, G2/M checkpoint, apoptosis, and p53 pathways (Fig. 6 A). IFN-stimulated genes (*Ifi203*, *Mndal*) and IFN transcriptional regulators (*Irf1*, *Irf2*, *Irf2bp2*) were consistently enriched in CreER *Stat3*^f/f^ HSCs compared with CreER HSCs (Fig. 6 B). These comparisons also confirmed the effective depletion of *Stat3* in CreER *Stat3*^f/f^ HSCs (Fig. 6 B). The CreER *Stat3*^f/f^ MPP cluster showed similar transcriptional responses, with enriched Myc target, IFN-γ, G2/M checkpoint, oxidative phosphorylation, and apoptosis pathways and upregulated IFN-associated genes and regulators (Fig. 6, C and D). qRT-PCR analysis confirmed the significant induction of IFN-associated genes in CreER *Stat3*^f/f^ BM cells compared with CreER BM or WT competitors from either mixed BM chimera background (Fig. 6 E). Collectively, these data suggest that the tamoxifen- and CreER-mediated deletion of *Stat3* skews HSCs towards myeloid and megakaryocyte/erythroid lineages and induces the cell-intrinsic activation of cell cycle–, stress-, apoptosis-, and IFN-associated transcriptional responses in HSCs and MPPs.

**Figure 6.**
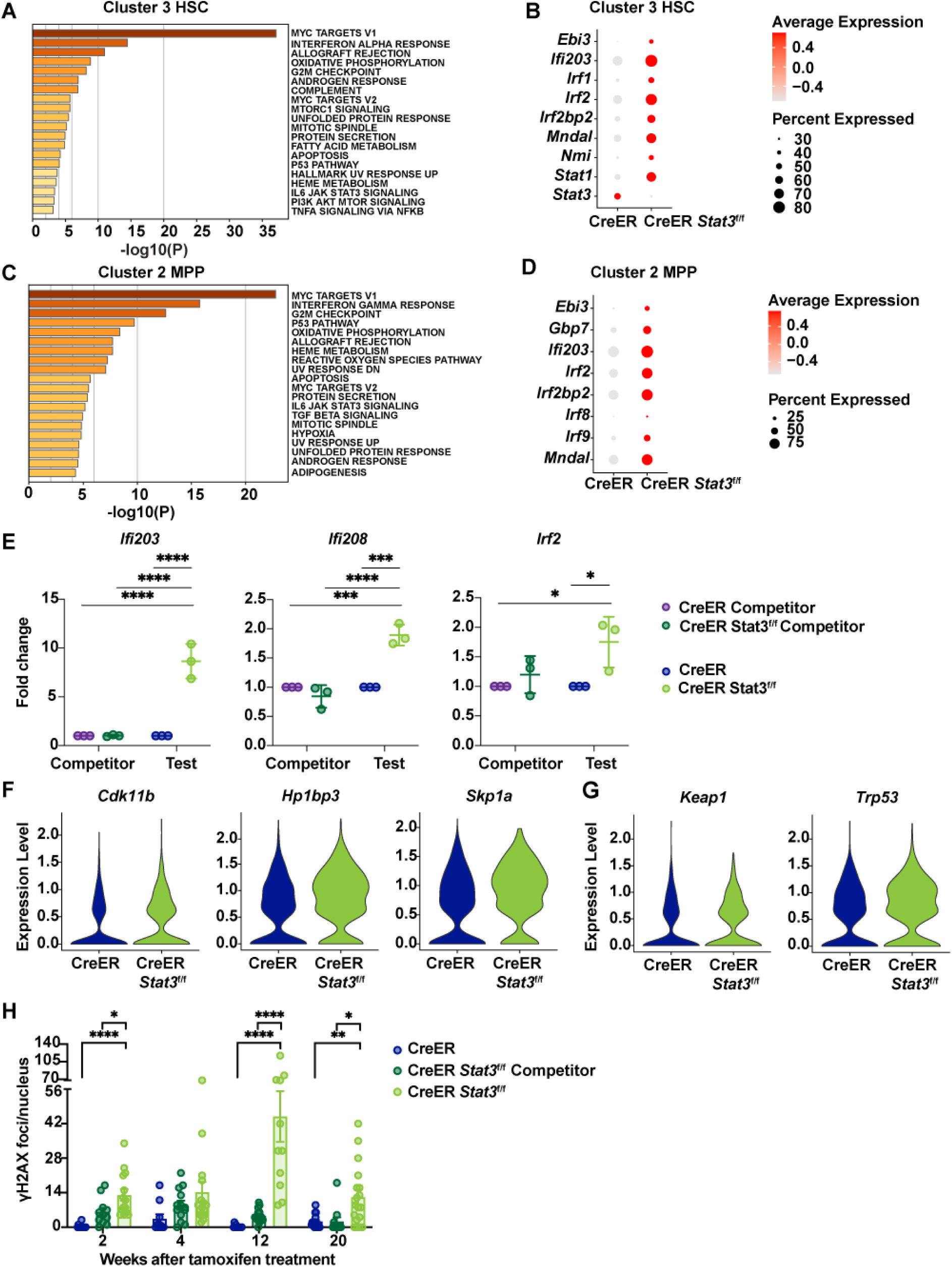
Enrichment of IFN signaling, cell cycle deregulation, and DNA damage in *Stat3*-deficient LSKs. (A-H) LSKs were purified from BM chimeric mice 8 weeks after tamoxifen treatment and processed for scRNA-seq analysis as indicated in Figure 5. (A) Pathway enrichment analysis of differentially expressed genes in cluster 3 (HSCs). The top 20 hallmark gene sets are shown for the comparison between CreER *Stat3*^f/f^ (*Stat3*-deficient) HSCs and CreER (*Stat3*-sufficient) HSCs. (B) Dot plot showing differentially expressed genes associated with the IFN transcriptional response in cluster 3 (HSCs) from the indicated experimental groups. (C) Pathway enrichment analysis of differentially expressed genes in cluster 2 (MPPs). The top 20 hallmark gene sets are shown for the comparison between CreER *Stat3*^f/f^ (*Stat3*-deficient) MPPs and CreER (*Stat3*-sufficient) MPPs. (D) Dot plot showing differentially expressed genes associated with the IFN transcriptional response in cluster 2 (MPPs) from the indicated experimental groups. (E) qRT-PCR analysis of IFN-inducible and -regulatory genes in total test (CreER or CreER *Stat3*^f/f^) and WT competitor BM cells from BM chimeric mice (n = 5 or 6 mice per group). (F) Violin plots showing the expression levels of genes associated with cell cycle regulation in cluster 3 (HSCs) from the indicated experimental groups. (G) Violin plots showing the expression levels of genes associated with stress response in cluster 3 (HSCs) from the indicated experimental groups. (H) Quantification of γH2AX foci. LSKs from the indicated experimental groups (n = 4-6 mice per group per timepoint) were purified and stained with DAPI (4’,6-diamidino-2-phenylindole) or with anti-γH2AX antibody prior to confocal imaging and analysis with Imaris software. Each data point represents the γH2AX foci count per nucleus for 1 field of view (n = 10-16 fields of view). In E and H, error bars indicate means ± SEMs. Statistical analyses were performed using 2-way ANOVA (E) or 1-way ANOVA (H) with the Tukey multiple comparison test. \**P* < 0.05; \*\**P* < 0.01; \*\*\**P* < 0.001; \*\*\*\**P* < 0.0001.

### *Stat3-*deficient HSPCs have an aberrant cell cycle and accumulate DNA damage

To examine the association of cell cycle–related transcriptional responses (e.g., those involving Myc targets, G2/M checkpoint, oxidative phosphorylation, and p53 pathways) with *Stat3* deficiency in HSCs and MPPs, we performed differential gene expression analyses using scRNA-seq data. Genes associated with cell cycle regulation (e.g., *Cdk11b, Hp1bp3, Skp1a*) were upregulated in CreER *Stat3*^f/f^ HSC and MPP clusters compared with the corresponding CreER subpopulations (Fig. 6 F and Fig. S4 A). Stress response genes (e.g., *Keap1, Trp53/p53)* also had higher expression in the CreER *Stat3*^f/f^ HSC cluster than in CreER HSCs (Fig. 6 G). These findings, in light of reduced cycling progenitors observed in scRNA-seq data, suggest that *Stat3*-deficient HSCs and MPPs have aberrant cell cycles.

Cell cycle deregulation or excessive HSC proliferation can lead to DNA damage, which in turn can activate intrinsic IFN responses (Bower et al., 2017; Brzostek-Racine et al., 2011, Hartlova et al., 2015, Yu et al., 2015). Therefore, to identify potential intrinsic sources of IFN induction in CreER *Stat3*^f/f^ LSKs, we quantified DNA damage in FACS-purified LSKs by measuring γH2AX foci over time (Fig. S4 B). CreER *Stat3*^f/f^ LSKs showed significant increases in γH2AX foci, particularly at 12 weeks after *Stat3* deletion (Fig. 6 H and Fig. S4 C), whereas CreER LSKs showed no significant differences in γH2AX foci over time. WT competitor LSKs from CreER *Stat3*^f/f^ BM chimeric mice showed a trend to accrue γH2AX at early time points after *Stat3* deletion, which is consistent with compensatory proliferation (Fig. 6 H). In addition, the DNA damage repair transcriptional pathway was enriched in CreER HSCs compared with WT competitors in the same background (Fig. S4 D). This transcriptional response was not induced strongly in CreER *Stat3*^f/f^ LSKs (Fig. S4 E), which suggests an impaired DNA damage response. Together, our results suggest that STAT3 is required to mediate the effective proliferation of HSPCs in response to a proliferative cue (i.e., tamoxifen), whereas the failure of this mechanism because of *Stat3* deficiency results in DNA damage, impaired DNA repair, and cell-intrinsic IFN signaling in LSKs.

### *Stat3* deletion in BM induces functional defects in HSCs

To formally evaluate whether *Stat3*-deficient HSCs are functionally defective, we utilized secondary BM transplantation assays. CD45.1^+^ congenic mice were injected with BM cells isolated from chimeric CreER or CreER *Stat3*^f/f^ mice 8 weeks after tamoxifen treatment (Fig. 7 A). Donor-derived CD45.2^+^ *Stat3*-deficent cells were significantly impaired in their ability to sustain hematopoiesis, as judged by an analysis of PB reconstitution in the secondary recipients (Fig. 7, B and C). Conversely, circulating CD45.1^+^CD45.2^+^ WT competitor cells were significantly increased in mice with CreER *Stat3*^f/f^ BM cells (Fig. 7 D). Twenty-four weeks after transplantation, total CD45.2^+^ cells and major CD45.2^+^ immune subsets were almost completely lost in the spleens of mice with CreER *Stat3*^f/f^ BM, whereas WT competitor subsets were increased (Fig. 7, E-H). Also, CreER *Stat3*^f/f^ BM failed to reconstitute BM LSK, LT-HSC, MPP-Mye, MPP-Ly, megakaryocyte-erythroid-biased MPP (MPP-MkE), MPP, GMP, CMP, common dendritic cell progenitor (CDP), CLP, and MEP populations (Fig. 8 and Fig. S5). Taken together, our results indicate that STAT3 is required for LT-HSC function in the absence of peripheral inflammation.

**Figure 7.**
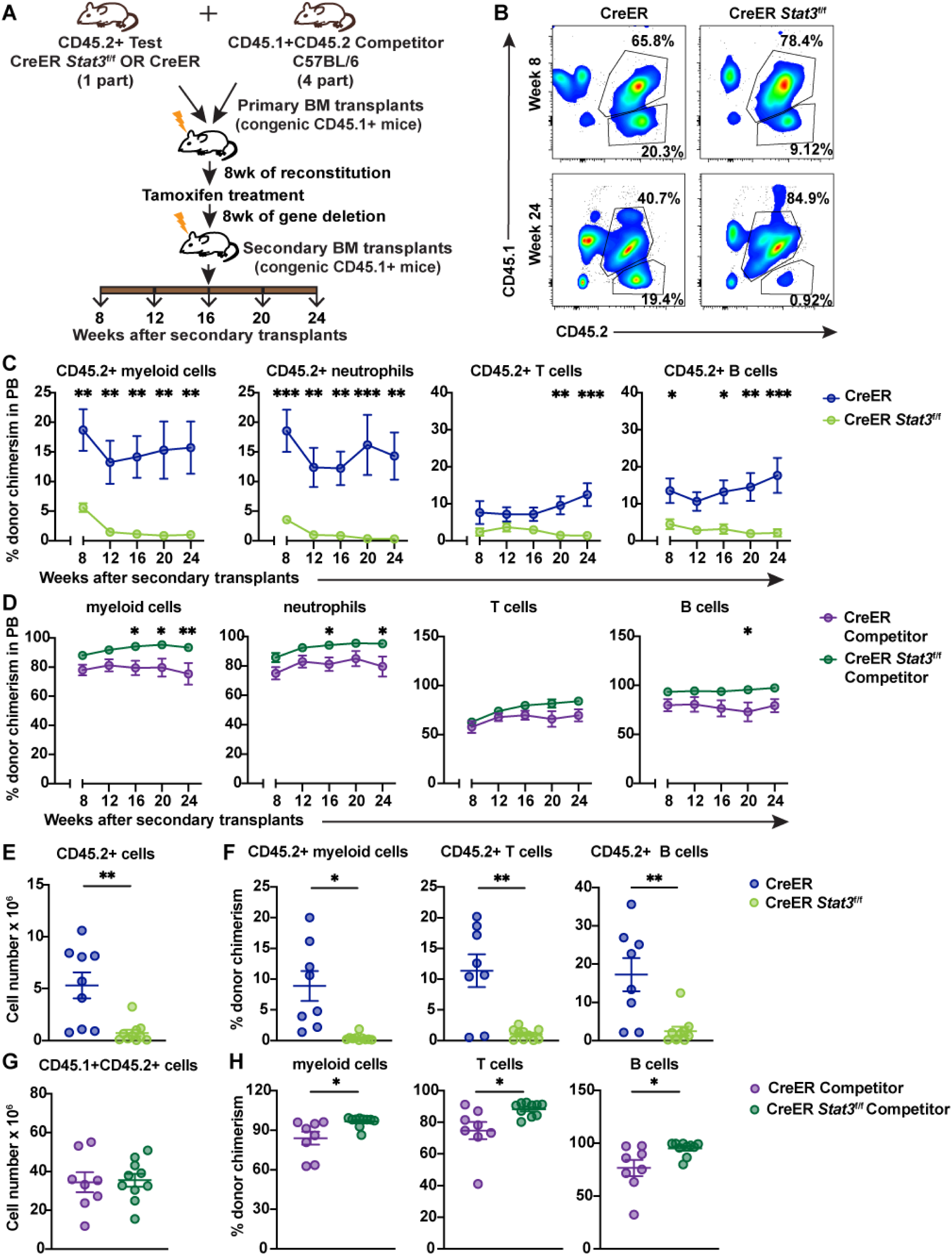
*Stat3*-deficient HSPCs fail to contribute to PB lineages after secondary BM transplantation. (A) A schematic diagram showing the experimental design for the secondary BM transplantation assays. (B) Representative flow cytometric plots showing proportions of CD45.2^+^ test and CD45.1^+^CD45.2^+^ WT competitor cells in PB after secondary BM transplantation. (C) Donor chimerism of CD45.2^+^ myeloid cells (CD11b^+^), neutrophils (CD11b^+^Ly6G^+^Ly6C^low^), T cells (CD3^+^), and B cells (B220^+^) in PB at the indicated times after secondary BM transplantation. (D) Donor chimerism of CD45.1^+^CD45.2^+^ myeloid cells (CD11b^+^), neutrophils (CD11b^+^Ly6G^+^Ly6C^low^), T cells (CD3^+^), and B cells (B220^+^) in PB at the indicated times. (E-H) Major immune cell lineages in the spleens were evaluated 24 weeks after secondary BM transplantation. (E) Absolute counts of CD45.2^+^ test cells. (F) Donor chimerism of CD45.2^+^ myeloid cells (CD11b^+^), T cells (CD3^+^), and B cells (B220^+^). (G) Absolute counts of CD45.1^+^CD45.2^+^ WT competitor cells. (H) Donor chimerism of CD45.1^+^CD45.2^+^ myeloid cells (CD11b^+^), T cells (CD3^+^), and B cells (B220^+^). Data in C-H are representative of 2 independent experiments (n = 8-10 mice per experimental group). Error bars indicate means ± SEMs. Statistical analyses were performed using 2-way ANOVA with the Sidak multiple comparison test (C, D) or using a 2-tailed unpaired Student *t*-test (E-H). \**P* < 0.05; \*\**P* < 0.01; \*\*\**P* < 0.001.

**Figure 8.**
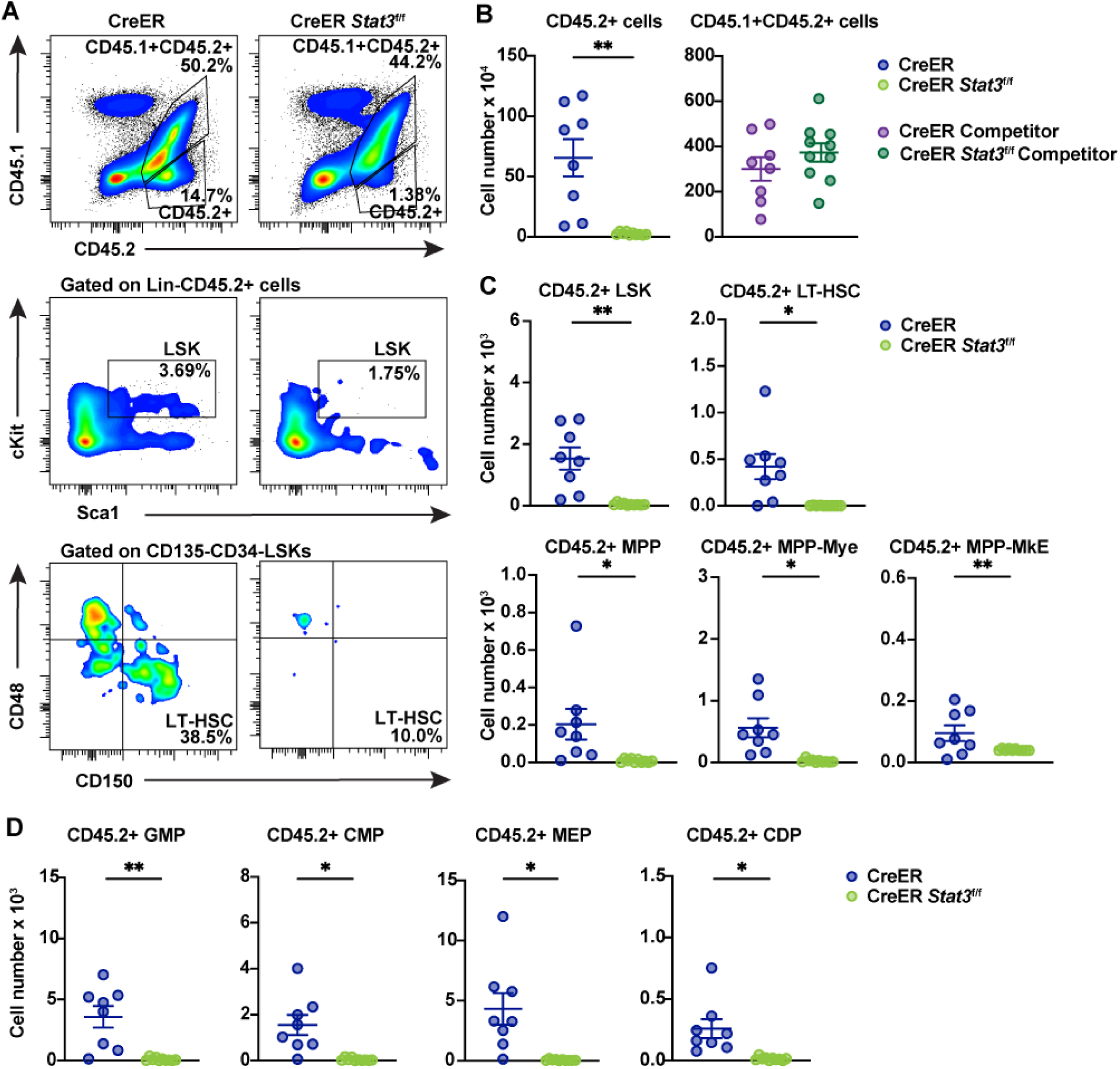
STAT3 is required to maintain LT-HSCs. BM progenitors were evaluated 24 weeks after secondary BM transplantation. (A) Representative flow cytometric plots showing proportions of CD45.2^+^ test and CD45.1^+^CD45.2^+^ WT competitor cells (top), LSKs (Lin^−^ ckit^+^Sca1^+^) (middle), and LT-HSCs (Lin^−^ckit^+^Sca1^+^CD135^−^CD34^−^CD48^−^CD150^+^) (bottom) in the indicated mice. (B) Absolute numbers of CD45.2^+^ test (left) and CD45.1^+^CD45.2^+^ WT competitor cells (right) in BM. (C) Absolute numbers of CD45.2^+^ test LSKs, LT-HSCs, MPPs (Lin^−^ckit^+^Sca1^+^CD135^−^CD48^−^CD150^−^), MPP-My (Lin^−^ckit^+^Sca1^+^CD135^−^CD48^+^CD150^−^), and MPP-MkE (Lin^−^ckit^+^Sca1^+^CD135^−^CD48^+^CD150^+^) in the indicated mice. (D) Absolute numbers of CD45.2^+^ test GMPs (Lin^−^CD127^−^ckit^+^Sca1^−^CD16/32^+^CD34^+^), CMPs (Lin^−^CD127^−^ckit^+^Sca1^−^ CD16/32^−^CD34^+^), MEPs (Lin^−^CD127^−^ckit^+^Sca1^−^CD16/32^−^CD34^−^), and CDPs (Lin^−^CD127^−^ ckit^low^CD115^+^CD135^+^) in the indicated mice. Data in B-D are representative of 2 independent experiments (n = 8-10 mice per experimental group). Error bars indicate means ± SEMs. Statistical analyses were performed using a 2-tailed unpaired Student *t*-test. \**P* < 0.05; \*\**P* < 0.01.

## Discussion

To address STAT3 function in HSPCs in non-inflammatory conditions, we employed a competitive mixed BM transplantation approach with CreER *Stat3*^f/f^ or CreER BM and WT competitor cells. This strategy enabled comparisons with prior work using mice transplanted with CreER *Stat3*^f/f^ BM in non-competitive conditions, which revealed key roles for STAT3 in HSPCs on an inflammatory background (Zhang et al., 2018). Our current use of mixed BM chimeras also allowed us to compare *Stat3*-deficient HSPCs with WT competitor cells in the same environment. Together, our results indicate that STAT3 protects HSCs and MPPs from aberrant DNA damage and IFN signaling in the context of a proliferative cue that is independent of external inflammatory signals. This protective response is required to maintain LT-HSCs and the effective production of mature immune lineages from the HSC pool.

Most HSCs are quiescent in homeostatic conditions but can transiently switch to a proliferative state during infection or other physiological stress events (Laurenti and Gottgens, 2018; Wilson et al., 2008). Several lines of evidence indicate that the CreER system induces a transient proliferative signal in the HSPC compartment. For example, we found in the present study that the proportions of CreER HSPCs in BM chimeric mice increase upon tamoxifen delivery and that tamoxifen stimulates the accrual of LT-HSCs in WT C57Bl/6J mice. Others have reported proliferative effects of tamoxifen in hematopoietic cells (Sánchez-Aguilera, 2016; Shehata et al., 2014). Therefore, we interpreted STAT3 function in the context of these findings. Our data indicate mouse models employing tamoxifen- and CreER-mediated gene deletion should be evaluated carefully, with appropriate comparisons between CreER cells and cells with CreER-mediated gene removal, as well as comparison of each cell type to WT cells in the same environment.

STAT3 governs G1/S transition and accelerates cell cycle progression in non-immune lineages and specific immune progenitor populations, particularly in response to cytokine cues (Fukada et al., 1998; Leslie et al., 2006; Sinibaldi et al., 2000; Zhang et al., 2010), yet whether STAT3 controls HSPC proliferation has been unresolved to date. Here, we found CreER *Stat3*^f/f^ LT-HSCs and progenitor populations failed to expand in BM chimeric mice in response to tamoxifen, suggesting a proliferative defect. Our scRNA-seq analysis revealed that several cell cycle and stress response transcriptional pathways were enhanced in CreER *Stat3*^f/f^ HSC and MPP clusters. In addition, expression of critical regulators of cell cycle and stress response pathways (Bai et al., 1996; Baird and Yamamoto, 2020; Dutta et al., 2014; Harris and Levine, 2005; Liu et al., 2017; Yu et al., 1998) were increased in CreER *Stat3*^f/f^ HSC and MPP populations. We also observed loss of cycling progenitors within the CreER *Stat3*^f/f^ HSPC compartment. Moreover, we found specific increases in WT competitor populations in CreER *Stat3*^f/f^ BM chimeric mice. Taken together, these results suggest *Stat3*-deficient HSPCs have a deregulated cell cycle leading to ineffective proliferation and compensatory growth of WT competitor cells in the same environment.

CreER *Stat3*^f/f^ LSKs also showed evidence of increased DNA damage compared with WT LSKs in the same chimeric background or with CreER LSKs from control cohorts. Persistent proliferative signals in HSCs can lead to DNA damage; moreover, cell cycle deregulation can damage nuclear DNA (Bower et al., 2017; Walter et al., 2015; Yanai and Beerman, 2020). Compared with WT competitors from the same background, CreER HSC and MPP clusters showed activation of DNA repair transcriptional responses, but CreER *Stat3*^f/f^ HSC or MPP clusters did not, which suggests that these populations had ineffective DNA repair. In addition, CreER *Stat3*^f/f^ HSCs or MPPs had impaired regulation of the DNA damage response gene *Trp53* (*p53*). Taken together, our data suggest that the deregulation of the cell cycle and DNA repair pathways in *Stat3*-deficient HSPCs leads to the accrual of DNA damage and ultimately the failure of *Stat3*-deficient HSCs.

Our scRNA-seq analysis also revealed that CreER *Stat3*^f/f^ HSC and MPP clusters had upregulated IFN-mediated transcriptional responses. DNA damage lesions induce specific signaling responses, including activation of signal transducers (e.g., STAT1), transcription factors (e.g., NF-κB, IRF3), and damage sensors (e.g., STING), that directly or indirectly activate cell-autonomous IFN signaling (Brzostek-Racine et al., 2011; Yu et al., 2015; Hartlova et al., 2015; Biechonski et al., 2017 and Li et al., 2016). In addition to having increased γH2AX, CreER *Stat3*^f/f^ BM cells had selective enrichment of IFN-response and -regulatory genes, which is consistent with the elevated IFN transcriptional responses due to DNA damage accumulation. Prior work from our laboratory indicates that STAT3 controls anti-inflammatory transcriptional responses in mature immune lineages by distinct mechanisms, including the suppression of NF-κB signaling in macrophages and the inhibition of autocrine IFN signal transduction in DCs (Chrisikos et al., 2022; Zhang et al., 2014). The results of our present study suggest that STAT3 can also inhibit inflammatory signaling by controlling appropriate cell cycling and DNA damage repair and restraining cell-intrinsic IFN signaling in HSPCs.

Type I and II IFNs have direct and indirect effects on hematopoiesis (Chen et al., 2015; Essers et al., 2009), including skewing differentiation towards myeloid and megakaryocyte lineages (MacNamara et al., 2011; Matatall et al., 2014; Schürch et al., 2014; Qin et al., 2017; Thongon et al., 2021). For instance, the intrinsic activation of IFN signaling promotes the megakaryocyte-skewed differentiation of telomere-dysfunctional HSCs (Thongon et al., 2021). In addition, IFN-γ–STAT1 signaling promotes megakaryocytic- and myeloid-biased differentiation in HSCs (Essers et al., 2011; Huang et al., 2007; Masumi et al., 2009; Matatall et al., 2014; Pietras et al., 2014; Rouyez et al., 2005). Moreover, HSCs from individuals with *JAK2V617F^+^* myeloproliferative neoplasms have increased IFN-α signaling that is marked by the elevated expression of *Stat1* and *Nmi* and accompanied by megakaryocyte differentiation bias (Tong et al., 2021). Our scRNA-seq data indicate that CreER *Stat3*^f/f^ LSKs have enhanced IFN transcriptional responses and have an increased accumulation of myeloid and megakaryocyte/erythroid subpopulations at the expense of lymphoid subpopulations. Furthermore, CreER *Stat3*^f/f^ HSC and MPP clusters showed elevated expression of myeloid- and megakaryocyte-related genes and transcription factors. These data suggest that the intrinsic activation of IFN signaling in *Stat3*-deficient HSCs and MPPs enhances the expression of myeloid- and megakaryocyte-related gene transcription and skews the differentiation of HSPCs towards myeloid and megakaryocyte progenitors.

We and others have found that *Stat3* deletion in hematopoietic cells induces myeloid-skewed hematopoiesis, yet these observations were made in the context of peripheral inflammation mediated by global hematopoietic *Stat3* deficiency (Mantel et al., 2012; Zhang et al., 2018). Our current study revealed evidence of myeloid- and megakaryocyte-biased hematopoiesis by CreER *Stat3*^f/f^ LSKs; however, we did not detect the enhanced production of peripheral myeloid populations in BM chimeric mice. In fact, myeloid subsets were rapidly depleted upon *Stat3* deletion in the non-inflammatory background of our model. The inability of *Stat3*-deficient myeloid progenitors to contribute to hematopoiesis in the absence of peripheral inflammation is consistent with prior work indicating that STAT3 has key roles in specific immune progenitors (Laouar et al., 2003; Zhang et al., 2010). By contrast, the expansion of myeloid populations from *Stat3*-deficient HSPCs in the context of systemic inflammation suggests that STAT3-independent mechanisms govern this response, which agrees with previous work indicating that additional pathways such as IFN–STAT1 or interleukin-1 signaling have important roles in myeloid-skewed hematopoiesis (King and Goodell, 2011; Matatall et al., 2016; Pietras et al., 2016).

Our secondary BM transplantation findings indicate that CreER *Stat3*^f/f^ HSCs cannot support hematopoiesis, which is consistent with the accrual of DNA damage in CreER *Stat3*^f/f^ HSPCs and the eventual depletion of *Stat3*-deficient cells in the HSC compartment (Biechonski et al., 2017; Kaschutnig et al., 2015; Moehrle et al., 2015). Thus, collectively our data suggest that STAT3 is essential to preventing catastrophic DNA damage in response to growth-promoting signals or as a consequence of aberrant proliferative cues and that these functions are required to preserve LT-HSCs. This information may help inform the design of novel synthetic lethal approaches to remove damaged or diseased HSPCs undergoing aberrant proliferation, which could be targeted for STAT3 inhibition. Investigating the feasibility of this approach will require new models to evaluate selective ablation of proliferating HSPCs with STAT3 inhibition while preserving quiescent HSCs. A full understanding of STAT3 activity in the HSPC compartment is needed to improve the therapeutic use of STAT3 inhibitors and prevent their unwanted off-target effects.

## Methods

### Mice

CreER *Stat3*^f/f^ mice were generated as described previously (Zhang et al., 2018). Age- and sex-matched CreER Gt(ROSA)26Sor^tm1(creERT2)Tyj^ animals (The Jackson Laboratory, Bar Harbor, ME) were used as controls unless mentioned otherwise (Ventura et al., 2007; Zhang et al., 2018). Congenic B6SJL-Ptprca Pepcb/boyj (CD45.1^+^) mice (The Jackson Laboratory) were used as BM transplant recipients (Janowska-Wieczorek et al., 2001; Zhang et al., 2018). CD45.1^+^CD45.2^+^ mice were generated by crossing congenic CD45.1^+^ mice with C57BL/6 (CD45.2^+^) mice. Animals were maintained in a specific pathogen–free facility at MD Anderson. All experimental procedures were performed in accordance with protocols approved by MD Anderson’s Institutional Animal Care and Use Committee.

### Competitive BM transplantation assays

BM cells were flushed from the femurs and tibias of donor mice into sterile DMEM containing 1% heat-inactivated fetal bovine serum. For primary mixed BM chimeras, BM cells from CreER or CreER *Stat3*^f/f^ (CD45.2^+^) mice were mixed with CD45.1^+^CD45.2^+^ wildtype (WT) competitor BM cells at a 1:4 ratio prior to transplantation. Eight-week-old congenic recipient CD45.1^+^ mice were lethally irradiated (920 rad) and intravenously injected with 2×10^6^ mixed BM cells. To evaluate reconstitution efficiency, we assessed PB every 4 weeks by flow cytometry.

For conditional *Stat3* deletion, beginning 8 weeks after BM transplantation, mixed BM chimeric mice were injected intraperitoneally with 2 mg of tamoxifen every other day for 1 week (3 treatments total). Hematopoietic activity was assessed by measuring PB chimerism every 4 weeks after tamoxifen treatment. For secondary transplantations, 2×10^6^ BM cells were isolated from mixed BM chimeric mice 8 weeks after tamoxifen treatment and transferred to lethally irradiated (920 rad) secondary CD45.1^+^ recipients. Hematopoietic activity was assessed by measuring PB chimerism every 4 weeks after secondary transplantation.

### Immune cell isolation and flow cytometry

For flow cytometry analysis, bone marrow (BM) chimeric mice were euthanized, and peripheral blood (PB), BM, spleen, and colon tissues were isolated. Single-cell suspensions of BM cells were prepared by flushing the cells from the bones and then performing cell dissociation as described previously (Zhang et al., 2018). To enrich lineage-negative (Lin^−^) hematopoietic stem and progenitor cells (HSPCs), BM cell suspensions were lysed in 1 mL of red blood cell lysis buffer (Tonbo Biosciences, San Diego, CA), washed with 1X phosphate-buffered saline (PBS), and incubated with a mixture of biotinylated antibodies directed against major hematopoietic lineage markers (Gr-1, CD11b, Ter119, CD3, and CD45R; 88-7774-75, Thermo Fisher Scientific, Waltham, MA) for 30 minutes. BM cells were subsequently washed, incubated with goat anti-rat IgG Microbeads (Miltenyi Biotec, Auburn, CA) for 30 minutes, and subjected to negative selection by magnetic-activated single cell sorting (Miltenyi Biotec) to purify Lin^−^ HSPCs as described previously (Zhang et al., 2018). Purified Lin^−^ HSPCs were washed with fluorescence-activated cell sorting (FACS) buffer (1X PBS supplemented with 2% fetal bovine serum and 2 mM EDTA) and incubated with the anti-mouse antibodies CD45.1 (A20), CD45.2 (104), CD127 (A7R34), cKit (2B8), Sca1 (D7), CD48 (HM48-1), CD16/32 (93), CD150 (TC1512F12.2), CD34 (Ram34), CD135 (A2F10), and CD115 (AFS98) (Table S4). Streptavidin eFluor 450 was added to exclude Lin^+^ BM cells. Lin^−^ HSPCs were incubated for 30 minutes, washed, and resuspended in FACS buffer prior to analysis by flow cytometry.

PB was collected in heparinized tubes, incubated with red blood cell lysis buffer for 10 minutes, and filtered through a 70-μM cell strainer. Single-cell suspensions were prepared from spleen by gently dissociating tissues through a 70-μM cell strainer with a 3-mL syringe plunger, using DMEM supplemented with 1% fetal bovine serum. Colonic lamina propria cells were isolated as described previously (Zhou et al., 2023). Blood, spleen, and colonic lamina propria cells were washed with PBS, blocked with anti-CD16/32 antibody (Tonbo Biosciences; 1:100) for 20 minutes, and then stained with fluorescently labeled antibodies for 30 minutes to detect major immune cell populations: CD45.1 (A20), CD45.2 (104), CD3 (17A2), B220 (RA3-6B2), CD11b (M1/70), Ly6G (IA8), Ly6C (HK1.4), CD11c (HL3), CD3 (17A2), MHC-II (M5/114.15.2), CD115 (AFS98), CD169 (3D6.112), CD64 (X54-5/7.1), CX3CR1 (goat IgG), CD62L (Mel-14), and CD103 (2E7) (Table S4). The stained cells were washed with PBS and resuspended in FACS buffer for flow cytometry analysis.

The frequencies, donor chimerism and absolute numbers of hematopoietic populations were determined. Donor chimerism was determined as the percentage of the target population within the combined CD45.1^+^, CD45.2^+^, and CD45.1^+^CD45.2^+^ cells of the same population. To calculate absolute numbers, precision counting beads (BioLegend, San Diego, CA) were added to each sample before flow cytometry analysis. Ghost dye (BV510, Tonbo Biosciences) was used for live/dead cell separation. Samples were run on an LSRFortessa X-20 cell analyzer (BD Biosciences, Franklin Lakes, NJ), and data were analyzed with FlowJo software (version 10.5.3).

### Multiplex cytokine/chemokine analysis

Protein lysates were generated from 1-cm proximal colon samples as described previously (Zhou et al., 2023). PB was obtained by cardiac puncture and allowed to clot for 1-2 hours at room temperature. Following centrifugation at 2000 rpm for 10 minutes (4°C), serum was removed and stored at −80°C until use. Colon lysates and serum samples were subjected to a bead-based multiplex assay (Cytokine & Chemokine 36-Plex Mouse ProcartaPlex Panel 1A, Invitrogen, Waltham, MA) performed on a Luminex 200 platform according to the manufacturer’s instructions. A 96-well plate assay was designed to analyze an array of 36 cytokines and chemokines (CCL2, CCL3, CCL4, CCL5, CCL7, CCL11, CXCL1, CXCL2, CXCL5, CXCL10, G-CSF, GM-CSF, IFN-α, IFN-γ, IL-1α, IL1-β, IL-2, IL-3, IL-4, IL-5, IL-6, IL-9, IL-10, IL12p70, IL-13, IL-15, IL-17A, IL-22, IL-23, IL-27, IL-28, IL-29, IL-31, LIF, M-CSF, and TNF-α). All samples were assayed in triplicate, and standards were measured in duplicate.

### Histology

Colon tissues were isolated, and a macroscopic assessment of inflammation was performed by measuring colon lengths. A 1.0-1.5 cm sample of the distal colon was washed with PBS, cut into 2 parts, and then fixed in 10% formalin. The fixed colon tissues were subjected to hematoxylin and eosin (H&E) staining by MD Anderson’s Research Histology Core Laboratory. Colon histology was assessed by trained pathologists using a previously established scoring system^2^ with a severity scale of 0 (healthy) to 4 (most disease).

### qRT-PCR

Total RNA was extracted from BM cells using Trizol (Life Technologies, Carlsbad, CA); 1 μg of RNA was reverse-transcribed to cDNA (Bio-Rad Real-time PCR system, Hercules, CA). *Stat3* deletion or differential gene expression between experimental groups was confirmed by performing qRT-PCR in a CFX real-time PCR machine (Bio-Rad) using 2X SYBR Select Master Mix (Sigma-Aldrich, St. Louis, MO) and gene-specific primers (Table S5). The expression of target genes was calculated using the 2^−ΔΔCT^ method with *Rpl13a* as an internal control (Pfaffl, 2000).

### Immunofluorescence

For the detection of γH2AX, 2000-3000 FACS-purified Lin^−^ckit^+^Sca1^+^ cells (LSKs; n = 4-6 mice per group) were subjected to centrifugation on glass slides using a Cytospin (Thermo Shandon, Pittsburgh, PA) at 5000 rpm for 10 minutes. The cells were fixed with ice-cold 100% methanol for 15 minutes at −20°C and blocked with PBS containing 5% normal goat serum and 0.3% Triton X-100 in a humidified container for 1 hour at 37°C. The cells were washed with Dulbecco’s PBS without calcium and magnesium (DPBS; Thermo Fisher Scientific) and then incubated with anti-γH2AX antibody (Cell Signaling Technology, Danvers, MA) at a 1:500 dilution in 1X PBS containing 1% bovine serum albumin and 0.3% Triton X-100 at 4°C overnight. The following day, the cells were washed 3 times with DPBS and incubated with secondary goat anti-rabbit antibody (Alexa Fluor 488; Thermo Fisher Scientific) at a 1:500 dilution in 1X PBS containing 1% bovine serum albumin and 0.3% Triton X-100 for 1-1.5 hours at room temperature. The cells were then washed, mounted with 10 µL of ProLong Diamond with DAPI (4’,6-diamidino-2-phenylindole; Molecular Probes, Eugene, OR), and incubated for 24-48 hours prior to analysis. Z-stack imaging of LSKs was performed on a Leica Sp8 confocal microscope equipped with a 63x objective lens (dimensions = 512 x 512; zoom factor = 6; pixel size = 60 nm x 60 nm) at MD Anderson’s Advanced Microscopy Core. Images were processed using Image J software (Schneider et al., 2012). A quantitative analysis of γH2AX foci per nucleus was performed using automated Imaris software (version 8.4.2, Bitplane, South Windsor, CT). Similar settings were used to enumerate foci across samples, and data were presented as the mean number of foci per nucleus.

### Single-cell RNA sequencing

We used fluorescence-activated cell sorting (FACS) to purify donor-derived test (CD45.2^+^) and competitor (CD45.1^+^CD45.2^+^) Lin^−^ckit^+^Sca1^+^ cells (LSKs) from mixed BM chimeric mice with CreER or CreER *Stat3*^f/f^ cells (n = 15 mice per experimental group) 8 weeks after tamoxifen treatment. Single-cell suspensions (1000 cells/µL) were submitted to MD Anderson’s Advanced Technology Genomics Core Facility. Cell viability was assessed, and samples were loaded onto a Chromium Single Cell Chip (10X Genomics, Pleasanton, CA) and processed for lysing and barcoding at a target capture rate of 10,000 single cells per sample. Barcoded libraries were pooled and sequenced using a NovaSeq 6000 100-cycle flow cell (Illumina, San Diego, CA) to an average depth of 100,000 reads per cell.

### Bioinformatics analysis of single-cell RNA sequencing data

Single-cell RNA sequencing (scRNA-seq) data were demultiplexed, and raw reads were aligned to the murine reference genome (mm10) using the CellRanger pipeline. CellRanger aggr was used to aggregate individual samples, and cell ranger gene expression matrices were further viewed and analyzed with the R package Seurat (version 3). Low-quality cells (<500 unique features) and cells with higher mitochondrial genome reads (>5%) were excluded from further analysis. The normalization of raw counts, dimensional reduction, and differential expression analyses were performed in Seurat. Pathway enrichment analysis of scRNA-seq data was performed using Metascape software (Zhou et al., 2019), and the top 20 hallmark gene sets were identified.

## Statistical analysis

Statistical analysis was performed using Prism (version 8.0, GraphPad, San Diego, CA). Significant differences were determined using an unpaired Student *t*-test or 1- or 2-way ANOVA with the Sidak or Tukey multiple comparison test.

## Online supplementary material

Fig. S1 shows hematopoietic activity in BM chimeric mice. Fig. S2 shows proportions of HSPCs in BM chimeric mice. Fig. S3 shows analysis of differential gene expression in LSKs from BM chimeric mice. Fig. S4 shows evaluation of the expression of genes associated with the cell cycle or with DNA damage repair response in LSKs. Fig. S5 shows evaluation of HSPCs in chimeric mice after secondary BM transplantation. Table S1 shows serum cytokine chemokine expression assessed by multiplex assay. Table S2 shows histological assessment of colon tissues. Table S3 shows colon length measurements of BM chimeric mice. Table S4 shows antibodies used for flow cytometry. Table S5 shows primers used for PCR analyses.

## Data Sharing Statement

The scRNA-seq data have been submitted to the NCBI GEO repository and are accessible through accession number (**GSE220466**).

## Acknowledgments

We thank Joseph Munch for his review and edits to this manuscript and Dr. Khandan Keyomarsi for advice and review of cell cycle data. This work was supported by grants from the Cancer Prevention and Research Institute of Texas (CPRIT) (Research Training award RP170067, R.L.B.) (Research Training award RP170067 RP210028, L.M.K. and E.M.P.), the National Institutes of Health (NIH) (R35 CA197566, F.G.G. and D.K.; R01AI133822 and R56AI109294-06, S.S.W.). S.C. is a Scholar of the Leukemia and Lymphoma Society. This work used MD Anderson’s Advanced Technology Genomics Core (supported by NIH 1S10OD024977-01 and NCI P30CA0166722), Research Histology Core Laboratory (supported by NCI P30CA0166722), Advanced Microscopy Core (supported by NIH S10RR029552), and Advanced Cytometry & Sorting Facility (supported by NCI P30CA0166722).

## Author Contributions

B.P. designed experiments, performed experiments, analyzed data, prepared data for presentation, and wrote the manuscript. Y.Z., R.L.B., Y.B.M., L.M.K., J.E.P., and E.M.P. assisted with sample collection and data analysis. D.K. assisted with scRNA-seq studies and analyzed data. F.M. curated scRNA-seq data, analyzed the results, and prepared data for presentation. M.A.Z. and T.Z. assisted with confocal microscopy and Imaris analysis. M.G.R. and X.T. performed pathology assessments. K.C.-D. assisted with flow cytometry data analysis. F.G.G. provided advice on scRNA-seq data analyses. S.C. analyzed scRNA-seq data, performed validation, provided advice on data interpretation and presentation. S.S.W. conceptualized the study, designed experiments, procured funding, supervised the study, and wrote the manuscript.

## Conflict of Interest Disclosures

Not applicable.

## Notes

### Competing Interest Statement

The authors have declared no competing interest.

