## Supplementary data for "STAT3 protects HSCs from intrinsic interferon signaling and loss of long-term blood-forming activity"

^1^Department of Immunology, The University of Texas MD Anderson Cancer Center, Houston, TX, USA; ^2^MD Anderson Cancer Center UTHealth Graduate School of Biomedical Sciences, Houston, TX, USA; ^3^Molecular Biology Institute, University of California, Los Angeles, Los Angeles, CA, USA; ^4^Division of Rheumatology, Department of Internal Medicine, Michigan Medicine, University of Michigan, Ann Arbor, MI, USA; ^5^Department of Leukemia, The University of Texas MD Anderson Cancer Center, Houston, TX, USA; ^6^Herbert Irving Cancer Center and Department of Genetics and Development, Columbia University, New York, NY, USA; ^7^Department of Surgical Oncology, The University of Texas MD Anderson Cancer Center, Houston, TX, USA; ^8^Department of Translational Molecular Pathology, The University of Texas MD Anderson Cancer Center, Houston, TX, USA; ^9^Department of Stem Cell Transplantation and Hematopoietic Biology and Malignancy, The University of Texas MD Anderson Cancer Center, Houston, TX, USA; and ^10^Program for Innovative Microbiome and Translational Research (PRIME-TR), The University of Texas MD Anderson Cancer Center, Houston, TX, US

**Correspondence:**

Stephanie S. Watowich, PhD

Department of Immunology, Unit 902

The University of Texas MD Anderson Cancer Center

1515 Holcombe Blvd.

Houston, TX 77030

**Supplementary Figure legends**

**Figure S1. Hematopoietic activity in BM chimeric mice.** PB samples from BM chimeric mice with CD45.2^+^ CreER or CreER *Stat3*^f/f^ test and CD45.1^+^CD45.2^+^ WT competitor cells were analyzed 4 and 8 weeks after tamoxifen treatment. (A) Donor chimerism of CD45.2^+^ myeloid cells (CD11b^+^). (B) Donor chimerism of CD45.1^+^CD45.2^+^ myeloid cells (CD11b^+^). (C) Donor chimerism of CD45.1^+^CD45.2^+^ neutrophils (CD11b^+^Ly6G^+^Ly6C^low^), monocytes (CD11b^+^Ly6G^-^Ly6C^high^), T cells (CD3^+^), and B cells (B220^+^). (D-H) Major immune cell lineages in the spleens and colons of BM chimeric mice were evaluated 8 weeks after tamoxifen treatment. (D) Donor chimerism of CD45.2^+^ myeloid cells (CD11b^+^) in spleen. (E) Frequency of CD45.1^+^CD45.2^+^ WT competitor cells in spleen (n = 12 or 13 mice per group). (F) Donor chimerism of CD45.1^+^CD45.2^+^ myeloid cells (CD11b^+^), neutrophils (CD11b^+^Ly6G^+^Ly6C^low^), monocytes (CD11b^+^Ly6G^-^Ly6C^high^), T cells (CD3^+^), and B cells (B220^+^) in spleen. (G) Donor chimerism of CD45.1^+^CD45.2^+^ DC subsets (CD11b^+^CD11c^+^ and CD11b^-^CD11C^+^) in spleen. (H) Frequency of CD45.1^+^CD45.2^+^ WT competitor cells (left) and donor chimerism of CD45.1^+^CD45.2^+^ myeloid cells (CD11b^+^; right) in colon (n = 9 or 10 mice per group). Data are representative of 2 independent experiments (n = 26-29 mice per group, A-C) (n = 9 or 10 mice per group, D-H) and error bars indicate means ± SEMs. Statistical analyses were performed using 2-way ANOVA with the Tukey multiple comparison test (A-C) or 2-tailed unpaired Student *t*-test (D-H). **P* < 0.05; ***P* < 0.01; ****P* < 0.001; *****P* < 0.0001.

**Figure S2. Proportions of HSPCs in BM chimeric mice.** (A) Gating strategy for HSPC subsets (LSKs, LT-HSCs, MPPs, MPP-Ly, MPP-My, MPP-MkE, and CLPs) of Lin^-^ BM cells. (B) Gating strategy for committed progenitor populations (GMPs, CMPs, MEPs, and CDPs) of Lin^-^ BM cells. (C and D) HSPCs in the Lin^-^ BM subset in BM chimeric mice were evaluated 0, 4, and 8 weeks after tamoxifen treatment. (C) Donor chimerism of CD45.2^+^ MPP-MkE (Lin^-^ckit^+^Sca1^+^CD135^-^CD48^+^CD150^+^), MPP-Ly (Lin^-^ckit^+^Sca1^+^CD135^+^CD150^-^), CMP (Lin^-^CD127^-^ckit^+^Sca1^-^CD16/32^-^CD34^+^), CLP (Lin^-^ckit^-^Sca1^-^CD115^-^CD127^+^CD135^+^), and CDP (Lin^-^CD127^-^ckit^low^CD115^+^CD135^+^) populations. (D) Donor chimerism of the indicated CD45.1^+^CD45.2^+^ HSPC populations. In C and D, data are representative of 2 independent experiments (n = 12 or 13 mice per group). Error bars indicate means ± SEMs. Statistical analyses were performed using 2-way ANOVA with the Tukey multiple comparison test (C and D). **P* < 0.05; ***P* < 0.01; ****P* < 0.001; *****P* < 0.0001.

**Figure S3. Analysis of inflammation and differential gene expression in LSKs from BM chimeric mice.** CD45.2^+^ and CD45.1^+^CD45.2^+^ LSKs were isolated by FACS from BM chimeric mice 8 weeks after tamoxifen treatment and subjected to scRNA-seq analysis as indicated in Figure 5. (A and B) Confirmation of non-inflamed conditions in the BM chimeric mice whose LSKs were used for scRNA-seq. (A) Colon histology assessments after 8 weeks after tamoxifen treatment. (B) Colon lengths of BM chimeric mice at the indicated times after tamoxifen treatment. (C) Feature plots of selected cluster-defining genes in the integrated LSK compartment. (D) Violin plots showing the expression levels of genes associated with myeloid (*Elane, Mpo, S100a8,* and *S100a9*), lymphoid (*Ighm* and *Jchain*), and megakaryocytic (*Pf4*) lineages in cluster 2 (MPPs) in CreER and CreER *Stat3*^f/f^ LSKs.

**Figure S4. Evaluation of the expression of genes associated with the cell cycle or with DNA damage repair response in LSKs.** LSKs from BM chimeric mice were processed for scRNA-seq as indicated in Figure 5. (A) Violin plots showing the expression levels of genes associated with cell cycle regulation and stress response in cluster 2 (MPPs) in CreER and CreER *Stat3*^f/f^ LSKs. (B) A schematic diagram showing the experimental design for assessing γH2AX amounts in LSKs. (C) Representative immunofluorescence images of γH2AX staining in LSKs. CD45.2^+^ LSKs were purified from CreER or CreER *Stat3*^f/f^ BM chimeric mice; CD45.1^+^CD45.2^+^ WT competitor LSKs were purified from CreER *Stat3*^f/f^ BM chimeric mice (n = 4-6 mice per group per timepoint). LSKs were stained with DAPI (blue) and with anti-γH2AX antibody (green). Magnification 63x. Scale bars represent 5 μm. (D) Pathway enrichment analysis of differentially expressed genes in cluster 3 (HSCs). The top 20 hallmark gene sets were identified in comparisons between CD45.2^+^ CreER and CD45.1^+^CD45.2^+^ WT competitor cells in the same background. (E) Pathway enrichment analysis of differentially expressed genes in cluster 3 (HSCs). The top 20 hallmark gene sets were identified in comparisons between CD45.2^+^ CreER *Stat3*^f/f^ and CD45.1^+^CD45.2^+^ WT competitor cells in the same background.

**Figure S5. Evaluation of HSPCs in chimeric mice after secondary BM transplantation.** HSPC populations in the Lin^-^ BM compartment were evaluated 24 weeks after secondary BM transplantation, mice were generated as indicated in Figure 7. (A) Donor chimerism of the indicated CD45.2^+^ HSPCs. (B) Absolute numbers of CD45.1^+^CD45.2^+^ WT competitor HSPCs. (C) Donor chimerism of the indicated CD45.1^+^CD45.2^+^ HSPCs. Data are representative of 2 independent experiments (n = 8-10 mice per experimental group). Error bars indicate means ± SEMs. Statistical analyses were performed using a 2-tailed unpaired Student *t*-test. **P* < 0.05; ***P* < 0.01.

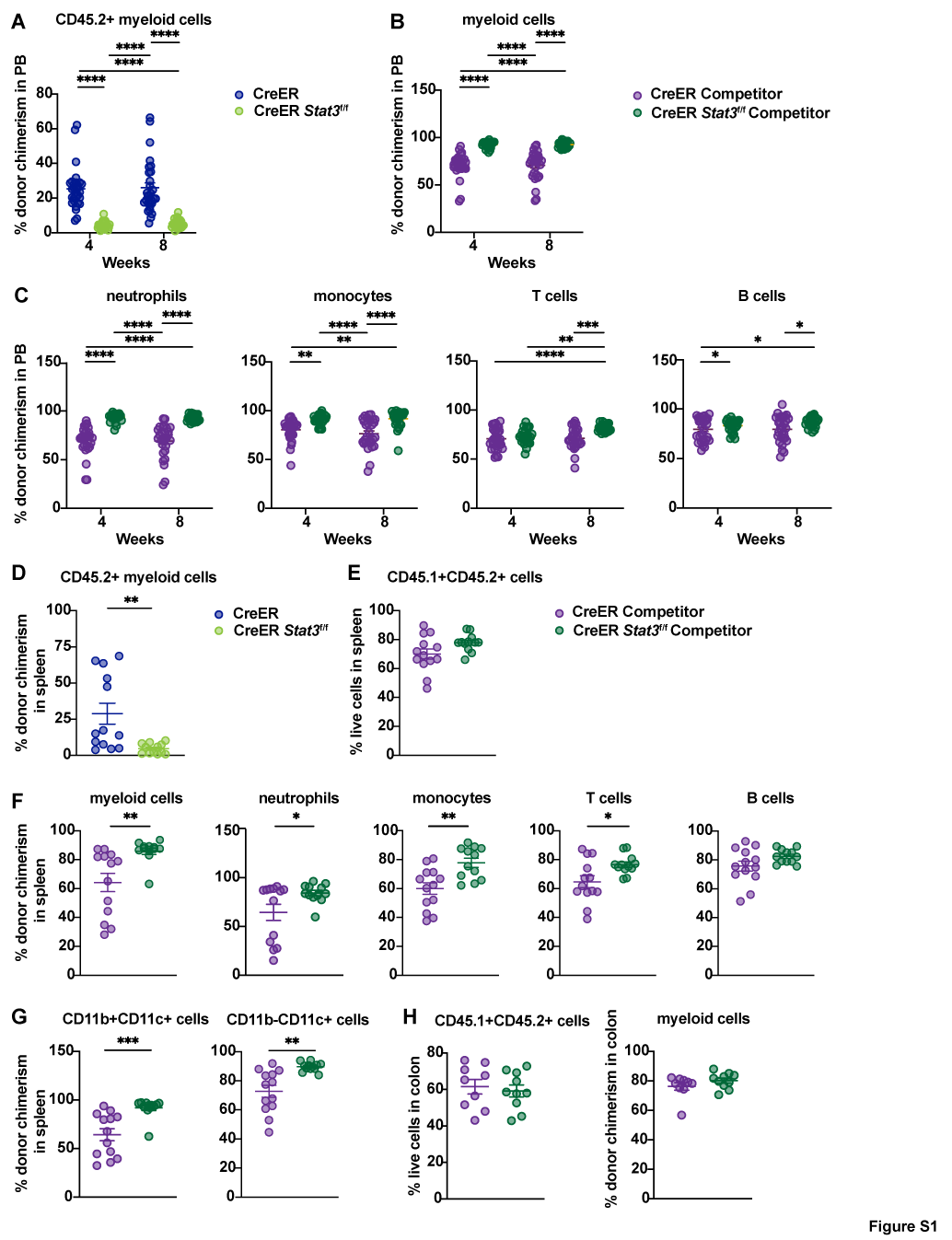

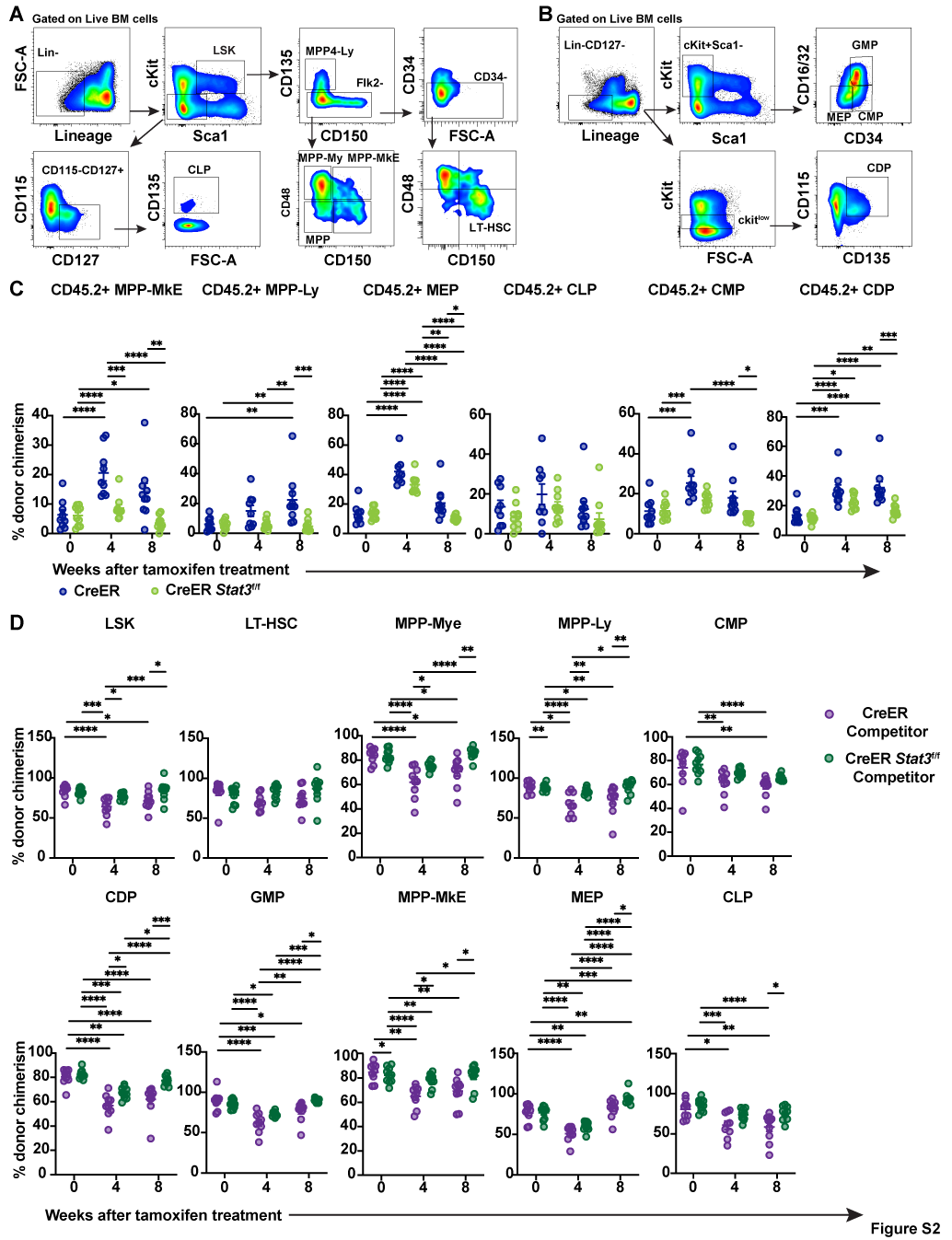

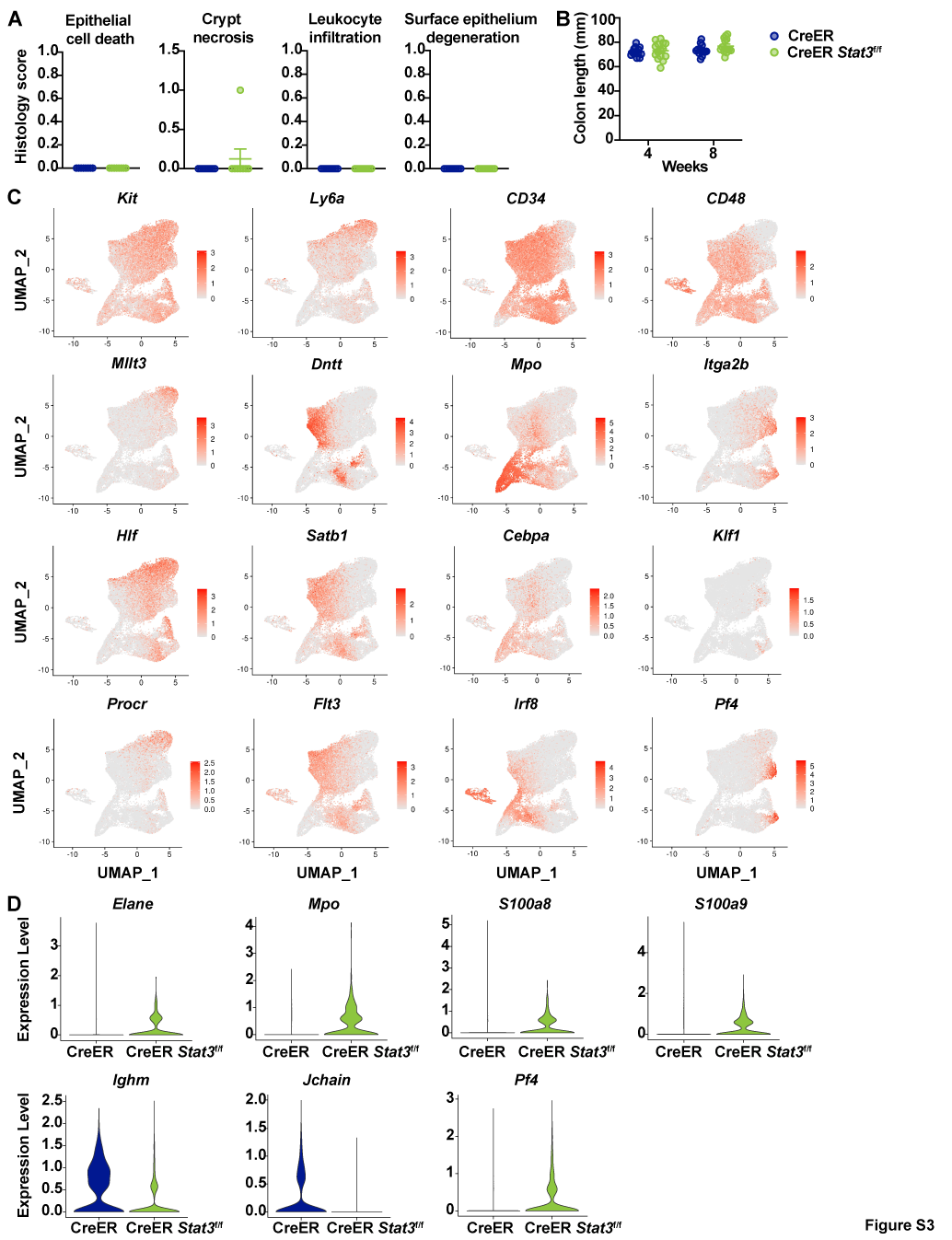

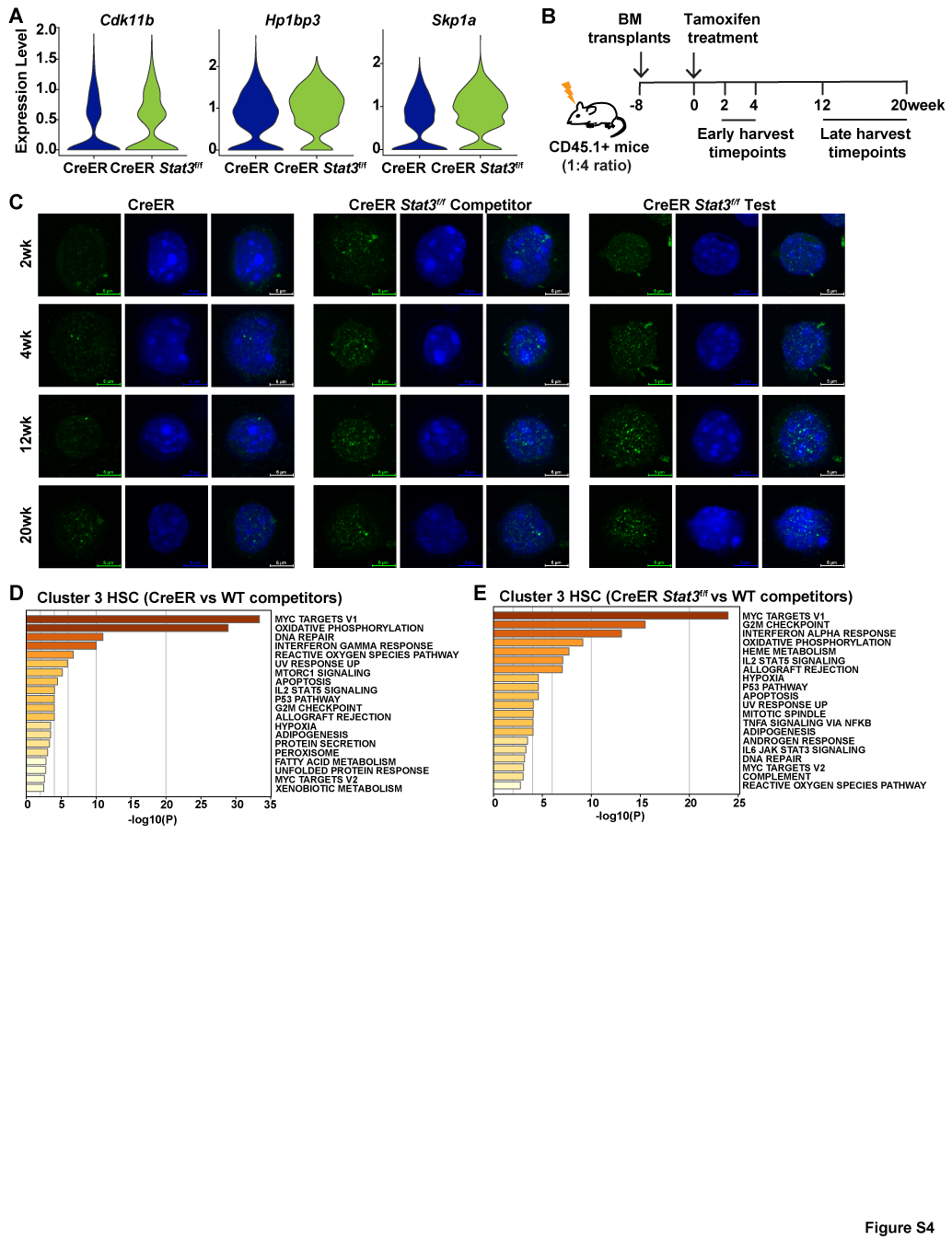

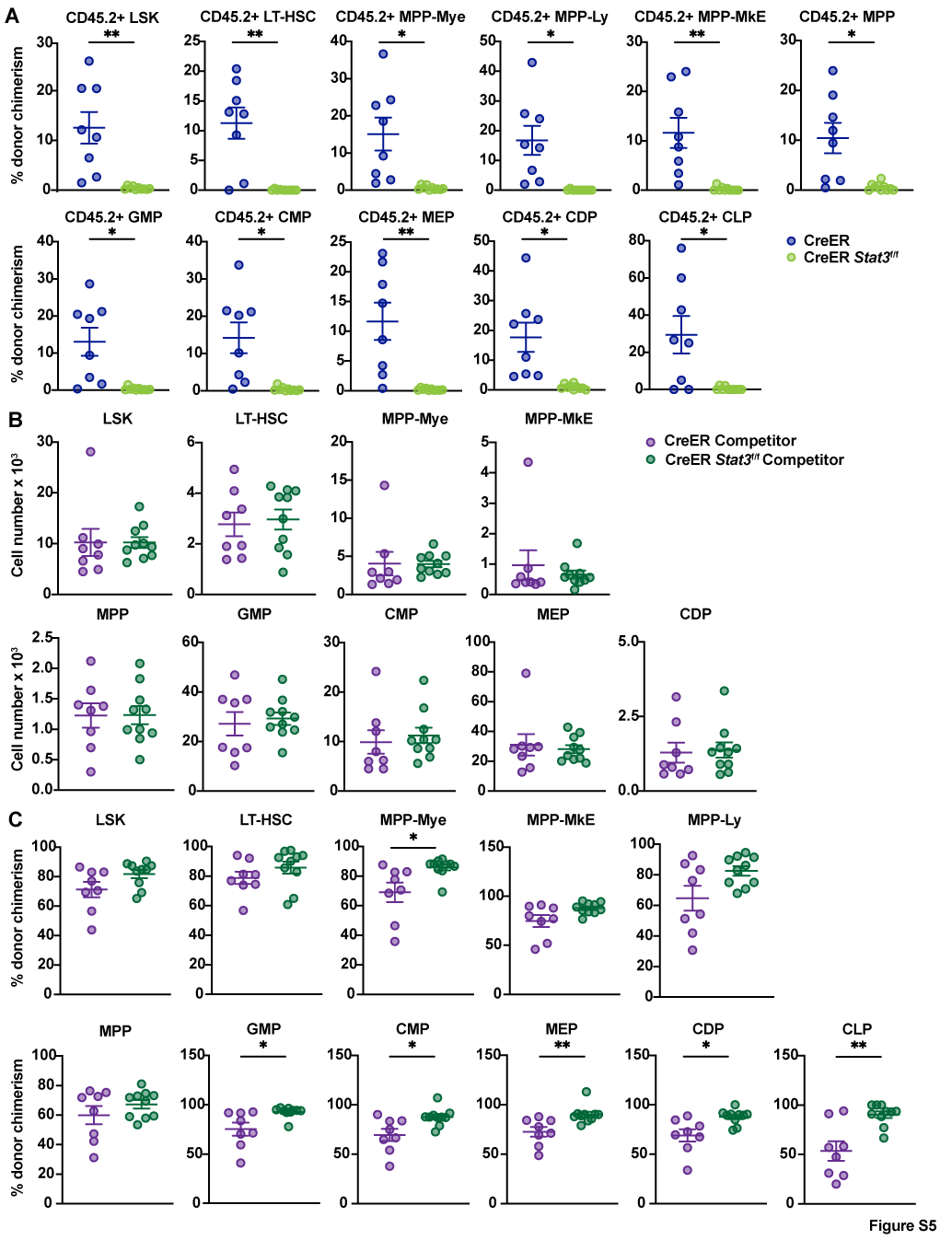

**Supplementary Tables**

**Table S1.** Serum cytokine chemokine expression assessed by multiplex assay

**Table S2.** Histological assessment of colon tissues

**Table S3.** Colon length measurements of BM chimeric mice.

**Table S4.** Antibodies used for flow cytometry

**Table S5.** Primers used for PCR analyses
